# The effects of warming on the stability of consumer-resource interactions

**DOI:** 10.1101/2023.09.27.559670

**Authors:** Alexis D. Synodinos, Arnaud Sentis, José M. Montoya, Bart Haegeman

## Abstract

Temperature regulates the physiology and behaviour of organisms. Thus, changing temperatures induce dynamics in species interactions. Considering that consumer-resource interactions underpin ecological communities, the impacts of warming on the stability of consumer-resource interactions have been extensively studied. However, a consensus among empirically determined warming-stability relationships and a clear understanding thereof is lacking. To investigate these systematically, we propose a simplified theoretical framework that can incorporate empirical data in three steps. First, we constrain stability to intrinsic oscillations to avoid comparing disparate stability notions. Second, we reduce complexity by utilising a one-dimensional stability metric. Third, we enable the direct comparison of all data by converting all thermal dependence parameterisations into a single function, with two parameters in the exponent determining its shape. The empirical data generate four different warming-stability relationships: stability increases, decreases, is hump-shaped or U-shaped with temperature. The diversity of warming-stability relationships, though partly attributable to context-dependence, is fundamentally caused by sensitivity to two factors: how the processes within the functional response are defined and the thermal dependence of carrying capacity. Consistency across studies regarding the former and acquiring more data on the latter should help uncover systematic patterns in the thermal dependence of stability in consumer-resource interactions.

## Introduction

Temperature affects the biology of organisms in many ways; it can cause changes in metabolic demands, foraging behaviour, ingestion rates and growth rates among other things (Pawar *et al*., 2016, Savage *et al*., 2004, Gillooly *et al*., 2001). Such organism- and population-level responses scale up to ecological communities and ecosystems via interspecific interactions (Montoya & Raffaelli, 2010, O’Connor *et al*., 2011, Thomas *et al*., 2004). Therefore, to predict how increasing ambient temperatures (IPCC, 2021) will alter communities, we first need to understand how interspecific interactions will respond to warming (Alvarez-Codesal *et al*., 2023, Bideault *et al*., 2021).

The dynamics of consumer-resource interactions are particularly important in this regard, as they regulate energy and material flows from primary producers towards higher trophic levels (Rip & McCann, 2011). Therefore, the impact of temperature change on stability — a composite aspect of dynamics (Donohue *et al*., 2013) — in consumer-resource dynamics has been extensively studied (e.g., Rall *et al*., 2010, Amarasekare, 2015, O’Connor *et al*., 2011, Fussmann *et al*., 2014, Amarasekare, 2019); however, the evidence appears inconclusive (Synodinos *et al*., 2021). In this study we seek to resolve this issue by producing a comprehensive overview of the warming-stability relationships which emerge from empirical data.

Theory postulates that stability properties can be deduced from aggregates of empirically measurable quantities (Yodzis & Innes, 1992, Synodinos *et al*., 2021). One such aggregate is consumer energetic efficiency, which quantifies the energy gained through consumption relative to that lost through metabolic demands (Yodzis & Innes, 1992). A second aggregate, interaction strength, quantifies the impact of the consumer on the resource, either in the short-term through the functional response (Rall *et al*., 2010) or in the long-term via the change in resource equilibrium due to the consumer (Berlow *et al*., 1999, 2004). Both aggregates are closely linked to various aspects of stability: intrinsic oscillations, consumer persistence and asymptotic resilience.

A reduction with warming in either energetic efficiency or interaction strength has been associated with increased stability via the dampening or elimination of oscillations (Vasseur & McCann, 2005, Rall *et al*., 2008). Lower energetic efficiency or weaker interaction strength due to warming have also been argued to increase the risk of consumer extinction due to starvation (Sentis *et al*., 2012, Rall *et al*., 2012, Alvarez-Codesal *et al*., 2023). However, the latter would imply the proximity of the dynamics to the consumer extinction boundary and, therefore, a loss of stability associated to consumer persistence. Hence, decreasing energetic efficiency or interaction strength, induced by warming, could simultaneously increase stability in one sense (weaker oscillations) and reduce it in another (consumer extinction).

To resolve this issue, we focused exclusively on oscillations as the property of stability for which we sought general warming-stability relationships. Intrinsic (i.e., not induced by demographic or environmental perturbations) oscillations of coupled consumer and resource populations manifest when self-regulation (i.e., intraspecific density dependence) in the consumer is small compared to the strength of interspecific interactions. In the absence of self-regulation and under certain conditions, such as enrichment (the paradox of enrichment Rosenzweig, 1971), consumers temporarily overshoot their equilibrium causing resource depletion, which is followed by a decline in consumers. These fluctuations in the consumer and resource populations can become self-sustained, leading to periodic oscillations. Such intrinsic oscillations are linked to the temporal variability in population densities, a property of the dynamics which has been often studied in relation to stability to perturbations (i.e., coefficient of variation, Donohue *et al*., 2013).

Johnson & Amarasekare (2015) demonstrated that the product of attack rate, handling time and carrying capacity quantifies the tendency of dynamics to oscillate periodically. The product of attack rate and handling time determines the saturation of the functional response. Hence, it could be viewed as a measure of predation efficiency (Yodzis & Innes, 1992); more efficient consumer species achieve functional response saturation at lower resource abundances. Carrying capacity, on the other hand, has long been viewed as a proxy for environmental conditions, e.g., enrichment (Rosenzweig, 1971). Thus, the stability metric aggregates information relating to the species’ interaction and the environmental conditions. In fact, it is one of two components of interaction strength (Synodinos *et al*., 2021). Therefore, one could consider this stability metric as a proxy for predation efficiency, enrichment or interaction strength. Significantly, for the purpose of our study, this is a one-dimensional metric which significantly reduces the complexity of the study of stability.

The systematic understanding of warming-stability relationships has also been hindered by an apparent lack of agreement in the empirically determined thermal dependencies of the relevant biological processes; attack rate, handling time and carrying capacity (Table 1). Certain studies report that vital rates follow a monotonic exponential relationship across a sufficiently wide temperature range (Rall *et al*., 2010, Fussmann *et al*., 2014), while others have shown that a non-monotonic pattern describes this temperature-dependence more accurately (Uiterwaal & DeLong, 2020, Uszko *et al*., 2017). In this case, after an exponential increase, the function peaks at a temperature referred to as optimal, before rapidly declining with further warming.

**Table 1:**
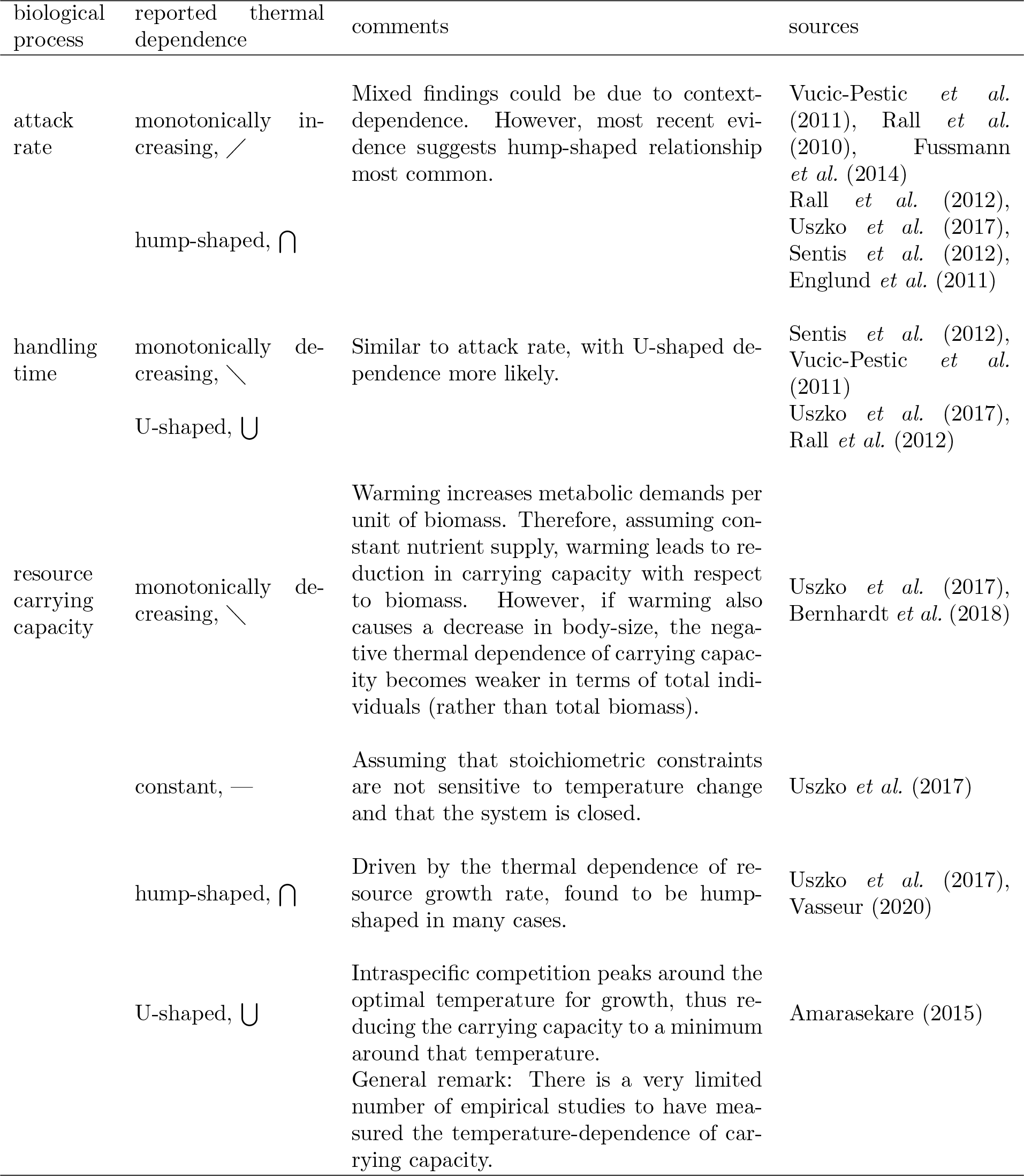
Evidence of the thermal dependencies of attack rate, handling time and carrying capacity from the literature.

Even though such differences in reported thermal dependencies of the various biological processes complicate the search of general patterns, a further issue has hindered the direct comparison between divergent empirical findings: thermal dependence data have been fitted with different types of functions across studies (e.g., Arrhenius equation, Gaussian function). To overcome this obstacle, we converted all empirically-determined parameterisations from the relevant literature into a single exponential form that can accommodate both monotonic and unimodal functions. We were, thus, able to directly compare empirical findings which were parameterised differently across study systems.

Thus, our approach was based on three steps: defining stability solely in terms of the tendency for intrinsic oscillations, using a one-dimensional stability metric and converting all thermal dependence parameterisations to a single function. Using this approach, we created a comprehensive overview of how the empirical evidence suggests warming will alter stability. Our results revealed four distinct warming-stability relationships. The difference in relationships was determined by three factors: the dichotomy between monotonic and non-monotonic thermal dependencies, the relative thermal sensitivities of attack rate and handling time and the — most often assumed rather than measured — thermal dependence of carrying capacity.

## Methods

We use the classical Rosenzweig-MacArthur model with a Holling type II functional response (eqs. 1 & 2), which will apply to a wide range of consumer-resource interactions (Jeschke *et al*., 2004). For the temperature dependence of organisms, we follow past studies in assuming a direct mapping of environmental temperatures to body temperature (Amarasekare, 2015, Vasseur & McCann, 2005) which best applies to ectotherm species. Using body temperature rather than ambient temperature as the temperature variable would overcome this constraint for other species without any impact on the qualitative patterns that we will describe. Further, we assume that resources have a wider thermal range of persistence than consumers.

### The model

The Rosenzweig-MacArthur model:

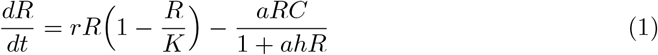

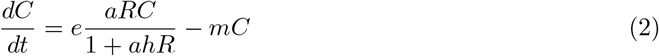

The equations describe the rate of change of resource, *R*, and consumer, *C*, biomass density ([*R*] = [*C*] = mass/area or [*R*] = [*C*] = mass/volume, where [*X*] denotes the units of variable *X*). The resource growth rate, *r* ([*r*]= 1/time), and carrying capacity, *K* ([*K*] = [*R*]) determine the logistic growth of the resource population. Consumers remove resource biomass through an increasing, saturating functional response (Holling type II). This is expressed as a function of attack rate, *a* (1*/*[*R*] *×* 1*/*time), and handling time, *h* ([*h*]=time). Consumer growth is proportional to the assimilated consumed biomass, with assimilation efficiency, *e* (unitless). Metabolic costs, *m* ([*m*] = 1/time), induce losses of consumer biomass. Below we briefly present the dynamics relevant to our analyses and provide only the necessary mathematical formulas; the full mathematical analysis relevant to our study is presented extensively in the supplementary material (Appendix S1).

### Model dynamics and stability metric

The model exhibits two qualitatively distinct dynamics (stationary regimes): stable equilibrium and stable limit cycle. In the former case, dynamics converge to the stable equilibrium either directly or via damped oscillations (App. S1, Fig. S1a, b). In a stable limit cycle, populations oscillate periodically around an unstable equilibrium (App. S1, Fig. S1c). A Hopf bifurcation occurs at the point where dynamics switch from stable equilibrium to stable limit cycle, though this transition is smooth rather than abrupt (App. S1, Fig. S2). We note here our assumption permeating our analyses; that dynamics have converged to the equilibrium or the limit cycle. This enables us to derive analytical expressions for the coexistence equilibria -whether stable or unstable -, for consumer and resource persistence and for the Hopf bifurcation (App. S1).

We focus on one aspect of stability: the tendency for intrinsic oscillations in population abundances. Johnson & Amarasekare (2015) proposed a single metric to quantify the tendency of dynamics to oscillate periodically in the Rosenzweig-MacArthur model (eq. 3).

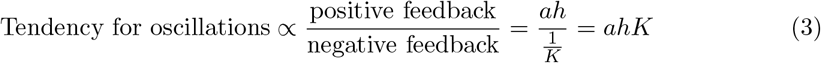

This metric revolves around two processes impacting *per capita* resource growth: a positive and a negative feedback. The positive feedback manifests via the functional response product *ah*. Once the functional response saturates, any additional resource abundance reduces the negative effect of consumption on per capita resource growth, promoting further resource growth. Consumers feed on the additional resources and overshoot their equilibrium. This causes the collapse in resource and then consumer abundances, thus inducing oscillations. The negative feedback occurs via resource self-regulation, which dampens fluctuations. In this model, this is expressed as the inverse of the carrying capacity (1*/K*). Therefore, *ahK* can be viewed as the ratio of positive to negative feedbacks in per capita resource growth (eq. 3).

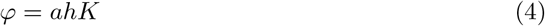

As has been previously shown, the Rosenzweig-MacArthur model’s dynamics can be simplified by grouping certain parameters together and deriving unitless aggregate parameters (Yodzis & Innes, 1992, Synodinos *et al*., 2021). One such aggregate, *φ*, is precisely the stability metric defined above (eq. 4, see also App. S1 for its derivation). As described in the introduction, *φ* encapsulates information on the trophic interaction (product *ah*) and the environmental conditions (*K*) such that it can be considered a proxy for interaction strength, enrichment or predation efficiency.

### Temperature-dependencies of parameters

We analyse the impacts of temperature on stability via the thermal dependence of *φ*, which is the product of three temperature-dependent parameters (eqn 4): attack rate, *a*, handling time, *h*, and carrying capacity, *K*. Our analysis is based on empirically determined thermal dependencies of these three parameters, as reported in the literature (Amarasekare, 2015, Vucic-Pestic *et al*., 2011, West & Post, 2016, Archer *et al*., 2019, Fussmann *et al*., 2014, Uszko *et al*., 2017, Binzer *et al*., 2016). Thus, we illustrate the different potential ways in which *φ* — and hence stability — depends on temperature.

In the literature, different functions have been used to parameterise the data on the thermal dependencies of *a, h* and *K* (e.g., Arrhenius, exponential quadratic with optimal temperature, etc., see Sup. inf. S2 for a comprehensive list). To simplify the analysis, improve comparability between studies and facilitate generalisations, we converted these parameterisations to a canonical form (eq. 5). This function, the general Gaussian, can produce both unimodal and monotonic thermal dependencies and represents a simple choice used previously in the literature (e.g., Amarasekare, 2015, Uszko *et al*., 2017). The transformations are based on a Taylor approximation (up to the second order) of the log-transformed temperature-dependent variable around the temperature *T*_0_ ≈ *T*. We present these transformations in the supplementary material (App. S2) and contend that they can provide a useful tool for comparing empirical findings.

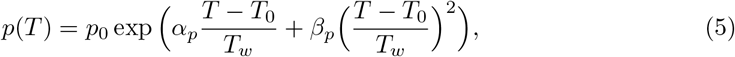

where

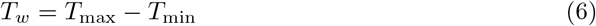

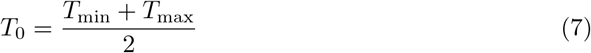

*T*_*w*_ (eq. 6) is the width of the temperature range of interest with *T*_0_ the mid-point of this temperature range (eq. 7). *T*_0_ should not be confused with the reference (*T*_ref_) or optimal (*T*_opt_) temperature in the original parameterisations. Finally, *p*_0_ is the parameter value at *T*_0_ and *α*_*p*_ and *β*_*p*_ are dimensionless constant parameters of the Gaussian function specific to *p*(*T*). We refer to these as linear and quadratic sensitivities, respectively. Depending on the relative magnitudes and the signs of *α*_*p*_ and *β*_*p*_, *p*(*T*) will attain different shapes (Fig. 1 and App. S2.1 for exact conditions).

**Figure 1.**
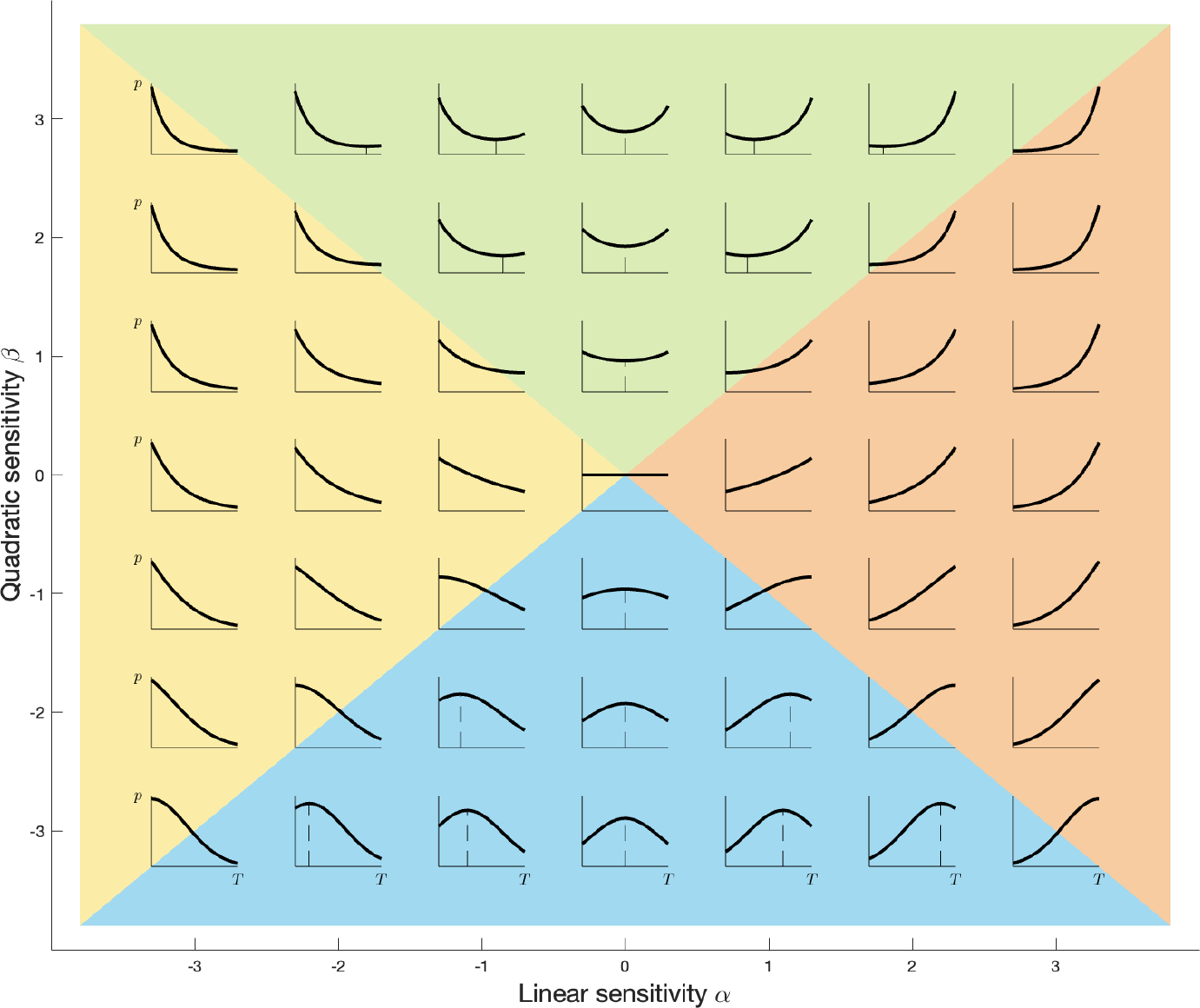
Graphical illustration of how the values of *α*_*p*_ (x-axis) and *β*_*p*_ (y-axis) determine the shape of the thermal dependence of *p*(*T*) (eq. 5). *α*_*p*_ and *β*_*p*_ determine the sensitivity of the exponential function (and hence the thermal dependence) on the linear and quadratic parts of the exponent, respectively. A relative dominance of *α*_*p*_ leads to a monotonic shape, while *β*_*p*_ relative dominance produces either hump- or U-shaped functions. Thus, four shapes emerge as defined by four regions, with smooth transitions between adjacent regions: monotonically increasing (red region), monotonically decreasing (yellow region), hump-shaped (blue region) or U-shaped (green region)

Our convention of using a single, general temperature parameterisation (eq. 5) simplifies and clarifies the analysis of the thermal sensitivity of stability. First, the transformed thermal dependence of individual parameters works well for all the original parameteristions we identified in the literature (App. S2, Fig. S3). Second, comparing like-for-like parameterisations across studies allows for the extraction of broader conclusions. Third, the stability metric, *φ*, becomes the product of three identical exponential functions. Mathematically, this simplifies things drastically: multiplying exponential functions amounts to adding the exponents. Given that *a, h* and *K* all share the same form, the values of *α*_*φ*_ and *β*_*φ*_ will be the sum of the respective *α*’s and *β*’s of *a, h* and *K*.

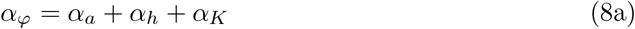

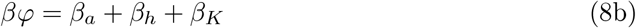

We note that our focus on the exponent — i.e., the shape of *φ* — provides the information on the relationship between temperature change and stability (i.e., increasing or decreasing the tendency for oscillations) which was the objective of our qualitative approach in the *α* − *β* plane. In the *α* − *β* plane, our findings are not sensitive to the value of the intercept *p*_0_.

## Results

### Four warming-stability outcomes

Warming-stability relationships derived from the literature were distributed across the *α* − *β* plane, indicating that our approach provided a good tool to encapsulate the available information and compare studies (Fig. 2). Significantly, our analysis revealed that empirical evidence fell into four distinct groupings for the thermal dependence of *φ*(*T*). Below we describe these patterns and look into the causes of this variation.

**Figure 2.**
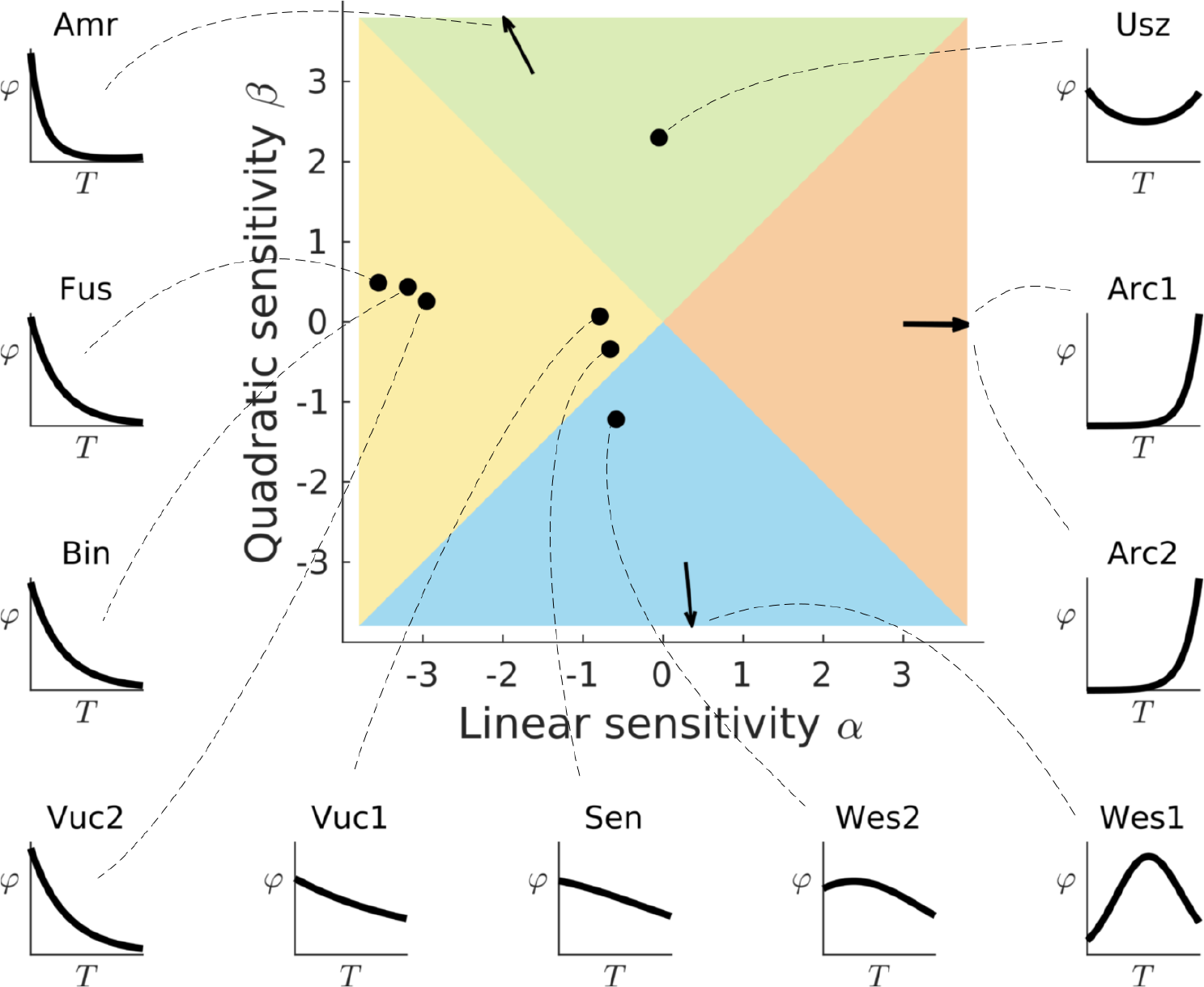
Computed values for *α*_*φ*_ and *β*_*φ*_ pairs based on empirical studies. The exact values for *α*_*φ*_ and *β*_*φ*_ are provided in App. 3. These are Amarasekare (2015), Archer *et al*. (2019), Binzer *et al*. (2016), Fussmann *et al*. (2014), Sentis *et al*. (2012), Uszko *et al*. (2017), Vucic-Pestic *et al*. (2011), West & Post (2016). In Archer *et al*. (2019), Arc1 and Arc2 correspond to two different years of measurement; in Vucic-Pestic *et al*. (2011), Vuc1 and Vuc2 correspond to two different prey types; in West & Post (2016), Wes1 and Wes2 correspond to two different predators. Studies in the yellow region have a monotonically decreasing *φ*(*T*), in the red region monotonically increasing *φ*(*T*), in the blue region hump-shaped *φ*(*T*) and in the green region U-shaped *φ*(*T*).

#### Scenario 1: stability increases with warming

Five studies yielded a monotonically decreasing *φ*(*T*), hence stability increased with temperature (Fig. 2 yellow region). These findings came from a meta-analysis of global invertebrate data (Fig. 2 Fus), from meta-analyses on soil invertebrate (for *K*) or unicellular organisms, invertebrate, ectotherm and endotherm vertebrates (for *a* and *h*) (Fig. 2 Bin), and from experiments of adult beetles feeding on flightless *Drosophila* adults (Fig. 2 Vuc1) or on beetle larvae (Fig. 2 Vuc2). The carrying capacity in both experiments (Vuc1, Vuc2) was not directly measured, but was based on literature sources, and decreased monotonically with temperature. Finally, a monotonically decreasing *φ*(*T*), which was close to the boundary with a hump-shaped *φ*(*T*), emerged from a predator-prey experiment of lady beetles feeding on a green peach aphid, (Fig. 2 Sen), with carrying capacity considered to be temperature independent.

#### Scenario 2: stability decreases with warming

A single study reported parameters producing an increasing monotonic thermal dependence for *φ*(*T*) (Fig. 2, red region: Arc1, Arc2); hence predicting a monotonic decrease in stability with warming. This study measured the thermal dependence of the interaction between two different freshwater predators, one sedentary (larvae of *L. riparia*, Arc1) and one mobile (larvae of *P. cingulatus*, Arc2), feeding on a single prey (blackfly larvae from the *Simuliidae* family). Carrying capacity, measured as the prey density without predators, increased with warming. The slope of *φ*(*T*) is quite steep, implying a very strong destabilising effect of warming.

#### Scenario 3: warming-stability relationship U-shaped

A single study produced a hump-shaped pattern for *φ*(*T*) (Fig. 2, blue region). Hence, stability was minimal around the optimum (U-shaped). The data points originate from experimental measurements of an algae (*Scendedesmus obliquus*) being consumed by two different *Daphnia* species, the small-bodied *D. ambigua* (Fig. 2 Wes1) and the large-bodied *D. pulicaria* (Fig. 2 Wes2).

#### Scenario 4: warming-stability relationship hump-shaped

Two studies gave rise to a U-shaped *φ*(*T*), implying that stability is maximised around the temperature optimum (Fig. 2, green region). The first sourced data from the literature on the thermal dependence of a cotton aphid (*Aphis gossypii*) and its parasite (*Aphidius matricatirae*) (Fig. 2, Amr). In this study, the minimum of *φ*(*T*) occurs close to the upper end of the temperature gradient, driven by a minimal carrying capacity at very high temperature. The second study was an experiment of *Daphnia hyalina* feeding on the green algae *Monoraphidium minutum* (Fig. 2 Usz). It is worth noting that carrying capacity was considered constant (i.e., no thermal dependence).

### Decomposition of warming-stability outcomes

Here we deconstruct the thermal dependence of *φ* into that of its three constituent biological parameters: attack rate, *a*, handling time, *h*, and carrying capacity, *K*. We, thus, illustrate how the four outcomes

#### Monotonic and non-monotonic thermal dependencies

Seven parameterisations produced a monotonic thermal dependence of *φ*. In six of these, the temperature-dependent (i.e., not constant) constituent parameters had a monotonic thermal dependence (Table 2, Arc1, Arc2, Bin, Fus, Vuc1 Vuc2). In the remaining case, attack rate was hump-shaped, handling time monotonic and carrying capacity constant (Table 2, Sen). Both exponents of handling time (*α*_*h*_, *β*_*h*_) were relatively large, with |*α*_*h*_| *> β*_*h*_ (see App. S2.4, Table S1 for all values). Attack rate had a quadratic exponent of similar magnitude but opposite sign to handling time, *β*_*a*_ *<* 0, and a relatively small linear exponent, *α*_*a*_ *>* 0. Thus, the sum of the two *β*s led to a smaller value than the sum of the *α*s (|*β*_*a*_ + *β*_*h*_| *<* |*α*_*a*_ + *α*_*h*_|). Hence, |*α*_*φ*_| *>* |*β*_*φ*_| and *φ*(*T*) was monotonic. The remaining four parameterisations yielded a non-monotonic *φ*(*T*) (Table 2, Amr, Usz, Wes1, Wes2). In these cases attack rate and handling time were non-monotonic, though carrying capacity was monotonic in one study (Table 2, Wes1, Wes2).

**Table 2:**
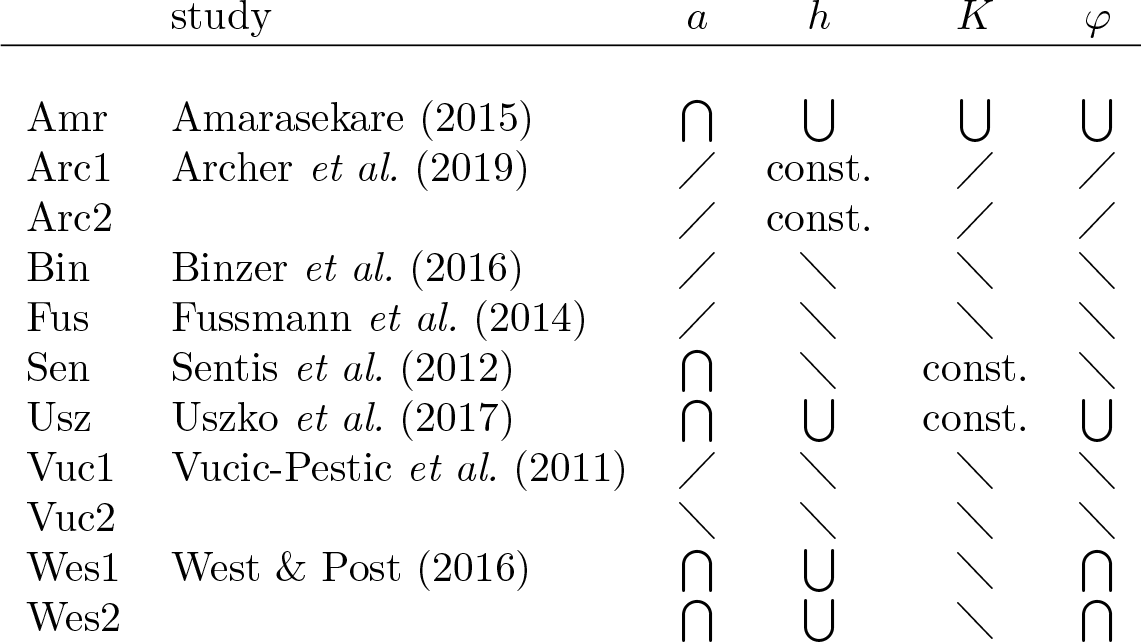
The thermal dependence shapes of attack rate, *a*, handling time, *h* and carrying capacity, *K*, and the resulting shaped in *φ*, in the studies used.

The split between monotonic and non-monotonic constituent parameters can provide a first clue for the shape of *φ*. However, the case of ‘Sen’ (Fig. 2, Table 2) illustrates it is not always robust. Next we look into a subtler difference that may provide an additional criterion in determining the shape of *φ* and, hence, warming-stability relationships.

#### Relative sensitivity of attack rate and handling time

In almost all parameterisations, *a* and *h* had the opposite thermal dependence (Table 2). *a* increased monotonically or was hump-shaped, while *h* decreased monotonically or was U-shaped, respectively. However, in each study, either *a* or *h* was more sensitive to temperature (i.e., they did not cancel each other out). This difference in sensitivity will have a significant impact on the warming-stability relationship, particularly in cases where the final parameter, *K*, has a weak thermal dependence. Two different studies of Daphnia species feeding on algae help to illustrate the importance of the relative thermal sensitivities of *a* and *h* (Fig. 3).

**Figure 3.**
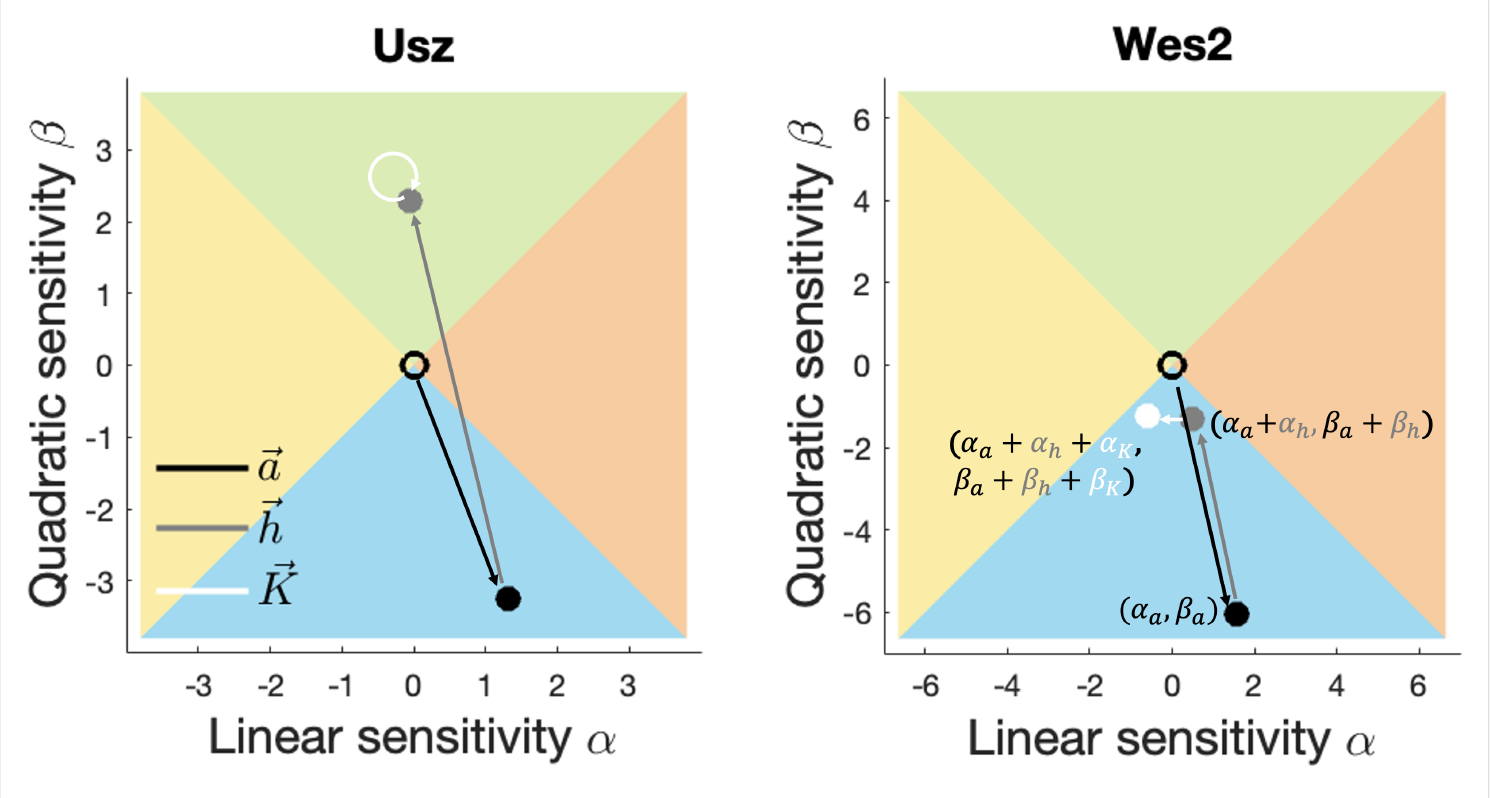
In these plots we illustrate how each individual parameter contributes to the outcome: starting from the origin (0, 0), each parameter can be represented as a vector whose projections on the *x*- and *y*-axis are determined by the *α* and *β* values, respectively. By consecutively adding the three vectors (one for each parameter), we end up with the position of *φ* in the plane. Here, attack rate, *a*, is depicted in black (arrow and point), handling time, *h*, in dark grey and carrying capacity, *K*, in white. In Usz, *K* has no thermal dependence (*K* = *constant*) and we included a loop-arrow to illustrate this. *a* is hump-shaped (*β*_*a*_ *<* 0) and *h* U-shaped (*β*_*h*_ *>* 0) with *β*_*h*_ *>* |*β*_*a*_|. Thus, *φ* is U-shaped (i.e., *β*_*φ*_ *>* 0). In Wes2, *a* and *h* are again hump-and U-shaped, respectively. However, in this case, *β*_*h*_ *<* |*β*_*a*_|. Moreover, the thermal dependence of *K* is both weak and monotonic, so its contribution in the *β* direction is minimal. Thus, *φ* emerges as hump-shaped.

In both cases, attack rate was hump-shaped and handling time U-shaped (Fig. 3 black and grey vectors, respectively). However, in the first study (Fig. 3 Usz), *a* had a weaker thermal sensitivity than *h*, so the addition of the two vectors landed *φ*(*T*) in the U-shaped region (green). Additionally, carrying capacity was assumed constant (Fig. 3, Usz white loop), thus not having any further impact on the thermal dependence of *φ*. In the second study (Fig. 3 Wes2), the opposite result was reported: *a* had a stronger thermal sensitivity than *h*. Combined with a weak thermal dependence of *K*, the thermal dependence of *φ* remained in the blue region, where *φ*(*T*) is hump-shaped.

#### Incomplete data on carrying capacity

Our approach shows that the contribution of each parameter to the sign and magnitude of *α*_*φ*_ and *β*_*φ*_ is equal (eq. 8a, 8b). Further, the thermal dependencies of *a* and *h* most often have opposite signs, and hence weaken each other. Therefore, the thermal dependence of carrying capacity could be significant in determining the thermal dependence of stability.

However, the thermal dependence of carrying capacity is rarely measured for the same study system as the functional response (*a, h*) parameters have been. From our data set, only one study did that (Wes1, Wes2 (West & Post, 2016)). For one study we used resource density observations (Archer *et al*., 2019), another used data from the literature sourced from different experiments on the same host-parasitoid pair (Amarasekare, 2015). The remaining parameterisations of carrying capacity were derived from a general (i.e., not species-specific) estimate of primary production (Vucic-Pestic *et al*., 2011), came from meta-analyses of species with no strong links to the consumer-resource pairs under consideration (Fussmann *et al*., 2014, Binzer *et al*., 2016) or were assumed constant (Sentis *et al*., 2012, Uszko *et al*., 2017).

### Sensitivity of warming-stability outcomes

For consumer-resource pairs close to the origin (i.e., where *α* = 0 and *β*=0), the thermal dependence of *φ* (and stability) will be sensitive to small changes in the individual parameters (e.g., due to measurement errors). Our analysis yielded three such cases (Fig. 2, Sen, Vuc1, Wes2), whose sensitivity arose because *a* and *h* had opposite thermal dependencies of similar magnitude and *K* did not have a strong thermal dependence (|*α*_*K*_|, |*β*_*K*_| ≲ 1).

## Discussion

The impacts of changing mean temperatures on consumer-resource interactions — and food webs by extension — have been studied through the thermal characterisation of specific biological rates (e.g., ingestion and respiration, Rall *et al*. (2010), Binzer *et al*. (2016)). These rates are used to parameterise consumer-resource models and, thus, derive the predicted dependence of trophic community dynamics on temperature (Vasseur & McCann, 2005, Rall *et al*., 2012, Kratina *et al*., 2022). A key aspect of dynamics which has been long studied is stability (Rosenzweig & MacArthur, 1963, Yodzis & Innes, 1992, Gilbert *et al*., 2014); however, we still lack a general expectation of how warming will alter the stability of consumer-resource interactions.

For one, the notion of stability has been applied to different properties of the dynamics Rall *et al*. (2010). A lack of stability has referred to consumer extinction (Sohlström *et al*., 2021, Daugaard *et al*., 2019), weaker asymptotic resilience (Gilbert *et al*., 2014) or increased tendency for intrinsic periodic oscillations (Johnson & Amarasekare, 2015, Kratina *et al*., 2022), though these three aspects of stability are not necessarily correlated (Yodzis & Innes, 1992, Uszko *et al*., 2017, Synodinos *et al*., 2021). Doubtless, this is not the first time that the complexity of the notion of stability has lead to confusion (Donohue *et al*., 2013, 2016, Grimm & Wissel, 1997).

To avoid such complications, we defined stability with respect to the tendency for intrinsic oscillations. These oscillations result from the intrinsic consumer-resource dynamics and should not be confused with oscillations induced by external perturbations. Further, we used a one-dimensional stability metric, the product of attack rate, handling time and carrying capacity (Johnson & Amarasekare, 2015) (eq. 3), which we termed *φ* (eq. 4). As a final step we converted the different functions used to parameterise the empirically determined thermal dependencies of attack rate, handling time and carrying capacity into a single, *canonical*, form (eq. 5). Thus, the shape of any empirically determined thermal dependence is dictated by two parameters of the canonical function, *α* and *β*, which determine the linear and quadratic sensitivity of the exponent, respectively.

Defining stability in terms of *φ* and using the canonical temperature dependence function helps illuminate why each study system yields a specific outcome. On one hand, the thermal dependence of *φ* encapsulates the combined effect of the three constituent parameters determining stability. On the other hand, the canonical form of temperature dependence maps the individual contribution of each parameter and the resulting thermal dependence of *φ* onto the *α*−*β* plane. Thus, our approach does not only provide a conclusive overview of the reported warming-stability relationships (Fig. 2), but it can further elucidate why each study system yields a given relationship through the interaction of attack rate, handling time and the carrying capacity (Fig. 3, Appendix S3.1, Fig. S5).

### No strict boundaries between warming-stability relationships

Our approach illustrates how warming-stability relationships fall into four groupings: stability can be monotonically increasing, monotonically decreasing, U-shaped or hump-shaped.

Though these outcomes appear distinct, they are not separated by some strict boundary. In particular, subtle differences in the measured or assumed thermal dependence of either attack rate, handling time or carrying capacity can cause a smooth transition into a different warming-stability outcome.

For instance, the parameter values reported in one study from our dataset were derived from a meta-analysis of global invertebrate data (Fussmann *et al*., 2014). The original study reported a stabilising effect of warming (Fig. 2, Fus). However, carrying capacity could arguably have a weaker thermal dependence either due to a concomitant reduction in body-size with warming (Bernhardt *et al*., 2018, Uszko *et al*., 2017). or by accounting for the uncertainty (i.e., variance) in the derivation of the mean parameter values of the meta-analysis. This uncertainty could also yield a slightly stronger thermal sensitivity of attack rate. In this case, warming would destabilise dynamics, producing the opposite warming-stability relationship compared to the original findings (App. S3, Fig. S6).

Changes in the choice of temperature range over which the analysis or experiments are conducted can also cause shifts between the outcomes. In our framework, the temperature range of interest was included within the general thermal dependence function (*T*_*w*_ in eq. 5). Altering the width of the temperature range can shift the warming-stability relationship in the *α* − *β* plane. In a specific example, Amarasekare (2015) reported that resource self-regulation (inverse of carrying capacity) peaked at 32°C, while attack rate and handling time had optima at 24°C and 22°C, respectively. Considering a temperature range wide enough to include the minimum in carrying capacity, results in a U-shaped *φ* (Fig. 2 Amr). However, if one constrains the temperature range to values below 32°C (e.g., due to experimental design limitations), carrying capacity would be an exponentially decreasing function. This would shift *φ* in the *α* − *β* plane into the region of monotonically decreasing thermal dependence and alter the warming-stability outcome (App. S3, Fig. S6).

These examples illustrate how the transition between different, opposite even, warming-stability relationships could occur smoothly, rather than as a result of a certain boundary in the system. We argue that such transitions are sensitive to subtle differences in the assumptions or measurements of the thermal dependencies of *a, h* or *K*. This is one reason that focusing on the individual empirically measured parameters has hindered generalisations. Instead, we need to consider their combined effects to be able to predict how temperature affects stability.

### Determinants of warming-stability relationships

Our analysis of the empirical evidence did not yield a single warming-stability outcome. Rather, studies were distributed across the two-dimensional plane, which represents the different shapes *φ* takes along the temperature gradient (Fig. 2). We identified three major contributing factors to this variation among findings.

First, there exists a dichotomy between the reported thermal dependence for both attack rate and handling time. Studies reporting a monotonic dependence for these parameters, yielded monotonic warming-stability outcomes (Table 2). Studies with non-monotonic attack rates and handling times produced non-monotonic warming-stability outcomes, with one exception (Table 2, Sen).

Overwhelming evidence suggests that attack rate and handling time have a hump- and U-shaped thermal dependence, respectively (Englund *et al*., 2011, Uiterwaal & DeLong, 2020). Nevertheless, divergence from this pattern has been reported (e.g., Vucic-Pestic *et al*. (2011), Rall *et al*. (2010), Archer *et al*. (2019)) and ascribed to context-dependence (Uiterwaal & DeLong, 2020). If one assumed that attack rate will tend to be hump-shaped and handling time U-shaped, would that yield fewer, a single even, warming-stability relationships? Probably not.

Given that attack rate and handling time have opposite thermal dependencies (increasing vs decreasing or hump-shaped vs U-shaped, respectively), the process with the strongest thermal dependence would shift the thermal dependence of stability accordingly in the *α* − *β* plane (e.g., Fig. 3). In one meta-analysis of ectotherms, attack rate had a steeper slope than handling time (Englund *et al*., 2011), whereas a more recent meta-analysis found evidence for the opposite (Uiterwaal & DeLong, 2020); handling time changed faster with temperature than attack rate. Thus, the second obstacle to simplifying generalisations arises.

The issue could originate in the different processes ascribed to handling time. In certain cases, handling time corresponds to the time required to ingest prey while in others it includes both the manipulation of prey and its ingestion (Jeschke *et al*., 2002). Interestingly, it has been shown that the former is more sensitive to temperature than the latter (Sentis *et al*., 2013). Thus, for predators limited only by ingestion, handling time could change faster than attack rate with temperature. On the other hand, predators limited by the manipulation and ingestion of the prey could have attack rates which change faster with temperature. A potential fix would be to break down the various processes represented within the functional response and to explicitly account for each one (Wootton *et al*., 2021, Sentis *et al*., 2013), or at minimum to explicitly define within each experiment what is being measured.

Third, the lack of data on the thermal dependence of carrying capacity constitutes a significant obstacle to uncovering systematic warming-stability relationships. Given how predictions can shift continuously (i.e., no discrete boundary) across the *α* − *β* plane, this lack of data could have profound effects. Therefore, the potential thermal dependence of carrying capacity requires careful treatment and ideally more measurements from the same study system for which the thermal dependence of the other parameters were established. Carrying capacity represents an aggregate of processes which ultimately constrain resource population growth (i.e., self-regulation), playing a significant role in the dynamics of populations and communities (Rosenzweig, 1971, Yodzis & Innes, 1992, Lemoine, 2019). Because it is hard to define as a specific biological quantity, empirical evidence for its thermal dependence remains scarce (but see Bernhardt *et al*. (2018)).

Measured as the resource equilibrium in the absence of consumers, carrying capacity has been shown to decrease with warming (O’Connor *et al*., 2009, Bernhardt *et al*., 2018, West & Post, 2016). An explanation has been attributed to the metabolic theory of ecology: given constant nutrient supply, per capita metabolic demands increase with temperature and cause the total population biomass to decline. Interestingly, if body-size decreases with temperature, then per capita metabolic demands increase less, dampening the effect of warming on the carrying capacity expressed in terms of individuals (Bernhardt *et al*., 2018).

Alternatively, the thermal dependence of carrying capacity could be hump-shaped. Based on a mathematical approach to the logistic growth function, the carrying capacity can be a linear function of the resource growth rate (Uszko *et al*., 2017, Vasseur, 2020). In this case, the carrying capacity would share the thermal dependence of the resource growth rate, which has been found to be hump-shaped (Amarasekare & Savage, 2012, Uszko *et al*., 2017). More realistically, carrying capacity emerges as an aggregate of nutrient supply, stoichiometry and metabolism among others (Uszko *et al*., 2017, Lemoine, 2019, Sentis *et al*., 2022). In this case, the hump-shaped thermal dependence remains the most likely outcome (Lemoine, 2019); however, temperature invariant or exponentially decreasing relationships have also been reported (Uszko *et al*., 2017).

Finally, carrying capacity could be U-shaped, as a result of stoichiometric constraints (Sentis *et al*., 2022) or from intra-specific competition. The latter has been shown to be hump-shaped, peaking at the same temperature as resource growth (Amarasekare, 2015). In the formulation of the logistic growth with intra-specific competition as self-regulation, carrying capacity would be expressed as the inverse of this competition. Therefore, carrying capacity would have a U-shaped thermal dependence.

Based on our discussion of the above scenarios, certain warming-stability relationships would appear more likely. If attack rate has a stronger sensitivity than handling time (i.e., combined manipulation and ingestion), their product would be hump-shaped (e.g., Fig. 3 Wes2). In this case, a hump-shaped carrying capacity would ensure a hump-shaped *φ*, so the warming-stability relationship would be U-shaped. This outcome would be still likely in the event of a monotonically decreasing carrying capacity (e.g., Fig. 3 Wes2). If, however, handling time (ingestion only) changes faster than attack rate with warming, then their product would be U-shaped (e.g., Fig. 3 Usz). A hump-shaped carrying capacity would counter this and weaken the overall thermal sensitivity of stability. Whether this would end up being hump-shaped or U-shaped, would depend on the details of the three thermal dependencies.

As we noted above, slight changes in assumptions or measurements can shift the thermal dependence of *φ*. Therefore, in the absence of sufficient data, it is hard to argue for the existence of general warming-stability relationships. Though it appears obvious, the best way to seek systematic patterns is to measure the thermal dependence of *φ* in multiple and well-defined experiments representing different systems.

### Caveats

Our approach of simplifying potential warming-stability outcomes invariably includes certain restrictive assumptions. Using *φ* reduces a complex problem to a one-dimensional metric. In certain cases, this could lead to a loss of accuracy as the consumer-resource pair approaches its thermal boundaries (Synodinos *et al*., 2021). Despite this, *φ* will give a good indication of the qualitative warming-stability relationship (Johnson & Amarasekare, 2015).

Relative to the thermal boundaries of the consumer-resource pair, we considered resources with a wider thermal niche than consumers. If this assumption failed, resource growth would become zero at temperatures for which consumers would not be physiologically limited. This would cause the abrupt loss of the resource (and consumer), however, our warming-stability predictions would not be affected if carrying capacity does not also become zero. If the carrying capacity also dropped to zero (Vasseur, 2020), then *φ* would rapidly decrease and stability — with respect to intrinsic oscillations — would increase prior to resource extinction.

## Conclusions

Understanding the temperature-dependence of consumer-resource dynamics is one of the building blocks for predicting how increasing ambient temperatures will impact food webs and ecological communities. Therefore, the stability of consumer-resource interactions and its thermal dependence represent a topic of much interest, however, with little consensus. We isolated a specific aspect of stability — the tendency for intrinsic oscillations — and simplified the analysis of empirically determined warming-stability relationships. Our approach mapped existing findings onto the thermal dependence of a single aggregate parameter, *φ*, summarising all empirical results in a simple plot. We further illustrated the usefulness of *φ* as a tool to decompose the warming-stability outcome into the individual contributions of attack rate, handling time and carrying capacity. The absence of obvious systematic impacts of warming on stability should not preclude the existence of such systematic effects. Though context-dependence did play a part, our analysis revealed how sensitive predictions can be to subtle assumptions or measurement errors. Well-defined experiments to characterise the thermal dependence of attack rate and handling time, and more measurements of the thermal sensitivity of carrying capacity should help reveal general warming-stability relationships. Our approach simplified a complex problem and provided a blueprint to improve our understanding. Ultimately, however, any resolution to the search for systematic warming-stability relationships rests on the existence and quality of sufficient empirical data.

