## Supplementary Information for "The effects of warming on the stability of consumer-resource interactions"

[**] Joint last authors

*Keywords*: climate change, predator-prey, population fluctuations, theory-data coupling, ectotherms, thermal biology

### Appendix S1: Rosenzweig-MacArthur model analysis

Here we expand on the mathematical analysis of the model. In the first section (S1.1), we derive the expressions for the Hopf bifurcation, the extinction boundaries of the resource and consumer, as well as the aggregate parameters which determine the dynamics. In the second section (S1.2), we illustrate graphically the different dynamical regimes and the transitions between them.

#### S1.1 Mathematical analysis

##### S1.1.1 The model

The Rosenzweig-MacArthur model:

$$\frac{dR}{dt} = rR\Big(1 - \frac{R}{K}\Big) - \frac{aRC}{1 + ahR} \tag{S1}$$

$$\frac{dC}{dt} = e\frac{aRC}{1 + ahR} - mC \tag{S2}$$

As noted in the main text, the equations describe the rate of change of resource, $R$, and consumer, $C$, biomass density ($[R] = [C] =$ mass/area or $[R] = [C] =$ mass/volume, where $[X]$ denotes the units of variable $X$). The resource growth rate, $r$ ($[r] = 1/\text{time}$), and carrying capacity, $K$ ($[K] = [R]$) determine the logistic growth of the resource population. Consumers remove resource biomass through an increasing, saturating functional response (Holling type II). This is expressed as a function of attack rate, $a$ ($[a] = 1/[R] \times 1/\text{time}$), and handling time, $h$ ($[h] = \text{time}$). Consumer growth is proportional to the assimilated consumed biomass, with assimilation efficiency, $e$ (unitless). Metabolic costs, $m$ ($[m] = 1/\text{time}$), induce losses of consumer biomass.

##### S1.1.2 Equilibria and stability

The dynamics are at equilibrium when the rate of change of both $R$ and $C$ is equal to zero. Setting both equations eq. (S1) and eq. (S2) to zero gives:

$$R_E = \frac{m}{a(e - mh)} \tag{S3}$$

$$C_E = er\frac{aK(e - mh) - m}{a^2K(e - mh)^2} \tag{S4}$$

The equilibria are positive if and only if eq. (S5) and eq. (S6) are satisfied:

$$e - mh > 0 \tag{S5}$$

$$aK(e - mh) - m > 0 \tag{S6}$$

If eq. (S6) (i.e. $C > 0$) holds, eq. (S5) is automatically satisfied and coexistence is guaranteed.

To determine the stability of the equilibria, we linearise the dynamics to derive the Jacobian matrix:

$$J = \begin{pmatrix} r(1 - \frac{2R}{K}) - \frac{aC}{(1+ahR)^2} & \frac{-aR}{1+ahR} \\ \frac{eaC}{(1+ahR)^2} & \frac{eaR}{1+ahR} - m \end{pmatrix} \tag{S7}$$

Then, we substitute $R$ and $C$ with the respective equilibrium values, $R_E$ and $C_E$. This will determine the stability of this equilibrium.

$$J_E = \begin{pmatrix} \frac{rm}{aeK(e-mh)}(ahK(e - mh) - e - mh) & \frac{-m}{e} \\ \frac{r}{aK}(aK(e - mh) - m)) & 0 \end{pmatrix} \tag{S8}$$

An equilibrium is stable if all eigenvalues of the Jacobian have negative real part. For a 2D dynamical system this condition is equivalent with the Jacobian having both a positive determinant and a negative trace (i.e., the sum of the diagonal elements). In eq. (S8) the determinant is always positive, so the equilibrium is stable if and only if the trace is negative, trace$(J) < 0$. The equilibrium is unstable if the trace is positive, trace$(J) > 0$. In this case, the dynamics oscillate around the unstable equilibrium (limit cycles). The point at which the dynamics switch from stable to limit cycles is termed a Hopf bifurcation and this corresponds to solution of trace$(J) = 0$:

$$ahK(e - mh) - e - mh = 0 \tag{S9}$$

##### S1.1.3 Parameter aggregation

Analyses of the Rosenzweig-MacArthur model have shown how unitless aggregate parameters (i.e., the grouping of parameters into an aggregate quantity) can be utilised to reduce the model's complexity (Yodzis & Innes, 1992, Vasseur & McCann, 2005, Synodinos *et al.*, 2021). Significantly, these aggregates are not simply a collection of other parameters. Rather, they correspond to biologically meaningful and empirically measurable quantities.

From eqs. (S5), (S6) and (S9), one can extract two such aggregate expressions, which we term $\rho$ and $\varphi$ (eqs. S10 & S11, respectively):

$$\varrho = \frac{e}{mh} \tag{S10}$$

$$\varphi = ahK \tag{S11}$$

$\varrho$ is the gain to loss ratio of assimilated maximal consumption ($e\frac{1}{h}$) over metabolic costs ($m$), often referred to as energetic efficiency. As Yodzis & Innes (1992) argued, this quantity is the maximum achievable energetic intake given only physiological constraints (i.e., when resource quantity is not limiting).

$\varphi$ is described at length in the main text. In brief, $\varphi$ includes two aspects of the community: the trophic interaction ($ah$) and the environmental conditions ($K$). The product $ah$ links the resource abundance to consumer growth. $K$ corresponds to the upper limit of the resource population at equilibrium (carrying capacity), which reflects the environmental conditions (e.g., enrichment, stoichiometric constraints, etc.).

We can now use these two aggregates, $\varrho$ and $\varphi$, to transform eqs. (S5), (S6) and (S9), respectively:

Resource equilibrium positive if: $$\varrho > 1 \tag{S12}$$

Consumer equilibrium positive if: $$\varphi(\varrho - 1) > 1 \tag{S13}$$

Hopf bifurcation, when: $$\varphi = \frac{\rho + 1}{\rho - 1} \tag{S14}$$

#### S1.2 Graphical illustration of dynamics

As we have shown with the existence of the Hopf bifurcation, the model has two distinct dynamical regimes: stable equilibrium and stable limit cycles. In the former case, dynamics either converge directly to the equilibrium (Fig. S1a, stable node) or they converge to the equilibrium via damped oscillations (Fig. S1b, stable focus). In the latter case, dynamics oscillate periodically around an unstable equilibrium (Fig. S1c, limit cycle).

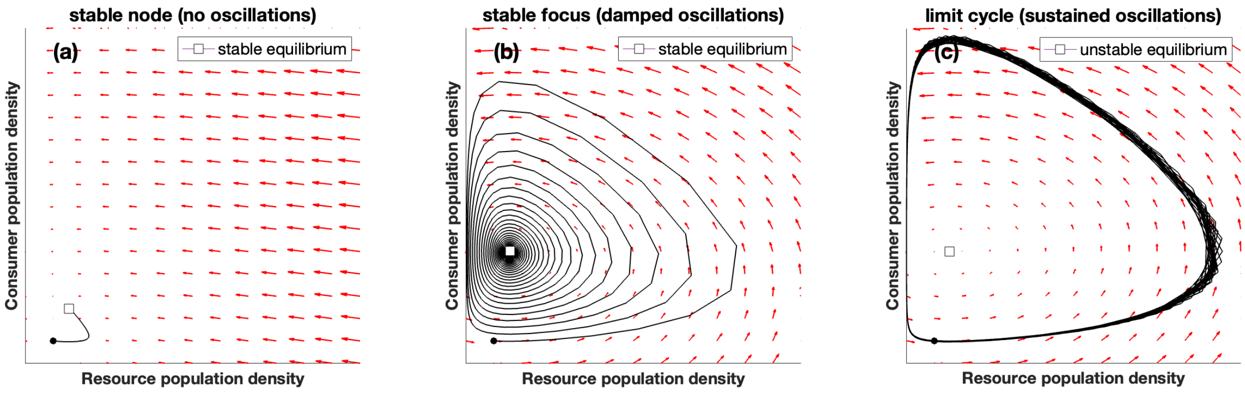


Figure S1: The phase portrait illustrates the trajectory (black curve) of the two population densities (resources on the x-axis, consumers on the y-axis) over time. (a) Population densities converge directly to the stable equilibrium. (b) Population densities initially oscillate. However, these oscillation converge towards the stable equilibrium, where populations eventually stabilise. (c) Population densities oscillate periodically as the dynamics converge to the stable limit cycle. The parameter values used for these plots were: $r = 0.5, a = 1, h = 0.1, m = 0.1, e = 0.65$ with $K = 0.3$ in (a), $K = 5$ in (b) and $K = 11$ in (c).

The dynamics in the phase portrait above correspond to certain temporal dynamics. Below, we illustrate that the transition from stable equilibrium to stable limit cycle occurs smoothly. In other words, as we approach the Hopf bifurcation, the damped oscillations persist for longer (Fig. S2), until the Hopf bifurcation is crossed and these oscillations become self-sustained.

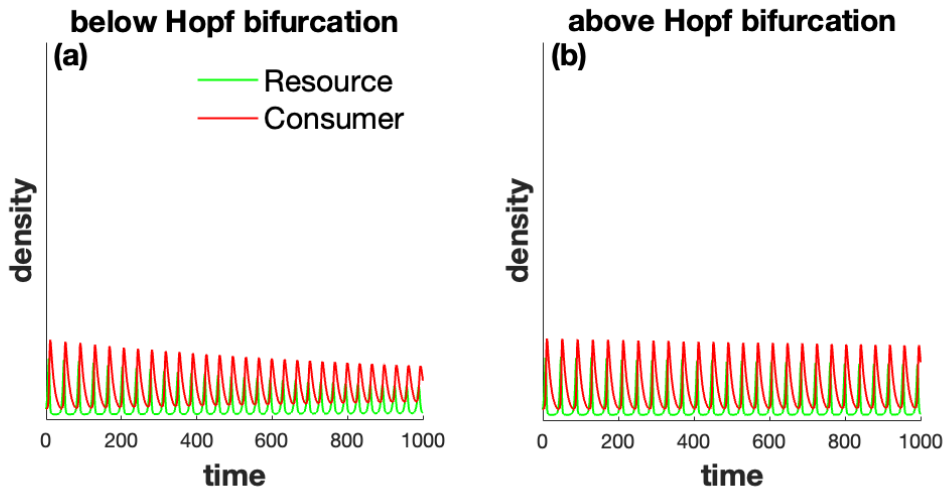


Figure S2: Timeseries of population densities close to the Hopf bifurcation. (a) Before dynamics cross the Hopf bifurcation, oscillations are damped. However, they persist for a very long time. (b) After crossing the Hopf bifurcation, the oscillations persist. The two timeseries illustrate that even though the Hopf bifurcation is determined by a strict condition (eq. S14), the transition between stable equilibrium and stable limit cycle occurs smoothly. The parameter values used for these plots were: $r = 0.5, a = 1, h = 0.1, m = 0.1, e = 0.65$ with $K = 9$ in (a) and $K = 10.5$ in (b). The Hopf bifurcation occurs at $K \approx 10.3$.

### Appendix S2: The *canonical* temperature-dependence function and related parameter values

The data we gathered from the literature consisted of parameterisations of the thermal dependence of attack rate, handling time and carrying capacity. These parameterisations utilised different functions (eqs. S25 - S29). We converted all the functions to a single form, referred to as *canonical* (eq. S15, the Gaussian). In this section we elaborate on these transformations in order clarify their details and make them readily available for anyone to use. First we present the canonical function and its parameters (S2.1). Following, we illustrate the conversion of the Arrhenius equation to the canonical form and remark on two equivalent forms of the Arrhenius equation which have been used interchangeably in the literature (S2.2). We, then, provide the parameter transformations required to convert different functions to the the canonical form (S2.3), before providing the exact parameter values in the original studies and the equivalent converted canonical functions (S2.4). Finally we illustrate the accuracy of the conversions graphically (S2.5).

#### S2.1 The *canonical* function

We selected the general Gaussian (eq. S15) as the function with which to represent all thermal dependencies. Our choice has significant advantages. First, it can reproduce all the thermal dependence forms reported in the literature. Second, its shape is determined by two parameters in the exponent, $\alpha$ and $\beta$ (eq. S15). This is critical because the stability metric is a product of three (temperature-dependent) parameters, $\varphi = ahK$. Therefore, the exponent of $\varphi$ will be the sum of the exponents of $a$, $h$ and $K$. On one hand, this constitutes a simple calculation and can be easily visualised in the $\alpha - \beta$ plane in terms of adding three vectors, corresponding to $(\alpha_a, \beta_a)$, $(\alpha_h, \beta_h)$ and $(\alpha_K, \beta_K)$. On the other hand, this allows one to trace any divergence in the resulting warming-stability relationships back to the thermal dependence of the individual parameters, $a$, $h$ and $K$. Below we elucidate the terms in the canonical function (eqs. S15−S17), illustrate the Taylor expansion with an example conversion of an Arrhenius equation into the canonical form and provide the parameter transformations to convert the functions from the literature (eqs. S25−S29).

The canonical function:

$$p(T) = p_0 \exp\Big(\alpha_p \frac{T - T_0}{T_w} + \beta_p \Big(\frac{T - T_0}{T_w}\Big)^2\Big), \tag{S15}$$

where

$$T_w = T_{\max} - T_{\min} \tag{S16}$$

$$T_0 = \frac{T_{\min} + T_{\max}}{2} \tag{S17}$$

$p_0$ is the parameter value at $T_0$ (eq. S15). $\alpha_p$ and $\beta_p$ are dimensionless constant parameters of the Gaussian function specific to $p(T)$. We refer to $\alpha$ and $\beta$ as linear and quadratic sensitivities, respectively. Below we will derive the conditions on $\alpha$ and $\beta$ which determine the shape of the canonical function (Fig. S3).

$T_w$ (eq. S16) is the width of the temperature range of interest. So, for an experiment measuring the thermal dependence of a biological rate between 5°C and 35°C, $T_w = 35 - 5 = 30$°C. $T_0$ is the mid-point of this temperature range (eq. S17). Continuing with the previous example, $T_0 = (35 + 5)/2 = 20$°C. If the original parameterisation was done with temperature in Kelvin, we add 273.15 K to the °C value in order to convert $T_0$ into Kelvin. Thus, $T_0 = 20$°C $= 293.15$ K. $T_0$ should not be confused with the reference ($T_{\text{ref}}$) or optimal ($T_{\text{opt}}$) temperature in the original parameterisations.

Here we derive the conditions for the canonical function to attain different types of temperature dependence (i.e., shapes). We introduce the rescaled temperature $\tau = \frac{T - T_0}{T_w}$, which varies over the interval $[-0.5, 0.5]$. The canonical function becomes

$$p(\tau) = p_0 \exp\big(\alpha_p \tau + \beta_p \tau^2\big) \tag{S18}$$

We know that:

- the temperature dependence always has an extremum, but this extremum can fall outside the valid temperature range $\tau \in [-0.5, 0.5]$;
- the sign of $\beta_p$ determines whether the extremum is a minimum ($\beta_p > 0$) or a maximum ($\beta_p < 0$);
- the extremum lies at $\tau = -\alpha_p / 2\beta_p$.

This leads to the following cases:

(a) minimum inside valid range: $\beta_p > 0$ and $|\alpha_p| < \beta_p$

(b) maximum inside valid range: $\beta_p < 0$ and $|\alpha_p| < -\beta_p$

(c) minimum left of valid range: $\beta_p > 0$ and $\alpha_p > \beta_p$

(d) minimum right of valid range: $\beta_p > 0$ and $-\alpha_p > \beta_p$

(e) maximum left of valid range: $\beta_p < 0$ and $-\alpha_p > -\beta_p$

(f) maximum right of valid range: $\beta_p < 0$ and $\alpha_p > -\beta_p$

Case (a) corresponds to a U-shaped and case (b) to a hump-shaped relationship. Cases (c) and (f) both generate an increasing relationship, and cases (d) and (e) both give a decreasing relationship. Hence, we obtain the following conditions:

U-shaped relationship: $\beta_p > 0$ and $|\alpha_p| < \beta_p$ (Fig. S3, green region)

hump-shaped relationship: $\beta_p < 0$ and $|\alpha_p| < -\beta_p$ (Fig. S3, blue region)

increasing relationship: $\alpha_p > 0$ and $\alpha_p > |\beta_p|$ (Fig. S3, red region)

decreasing relationship: $\alpha_p < 0$ and $-\alpha_p > |\beta_p|$ (Fig. S3, yellow region)

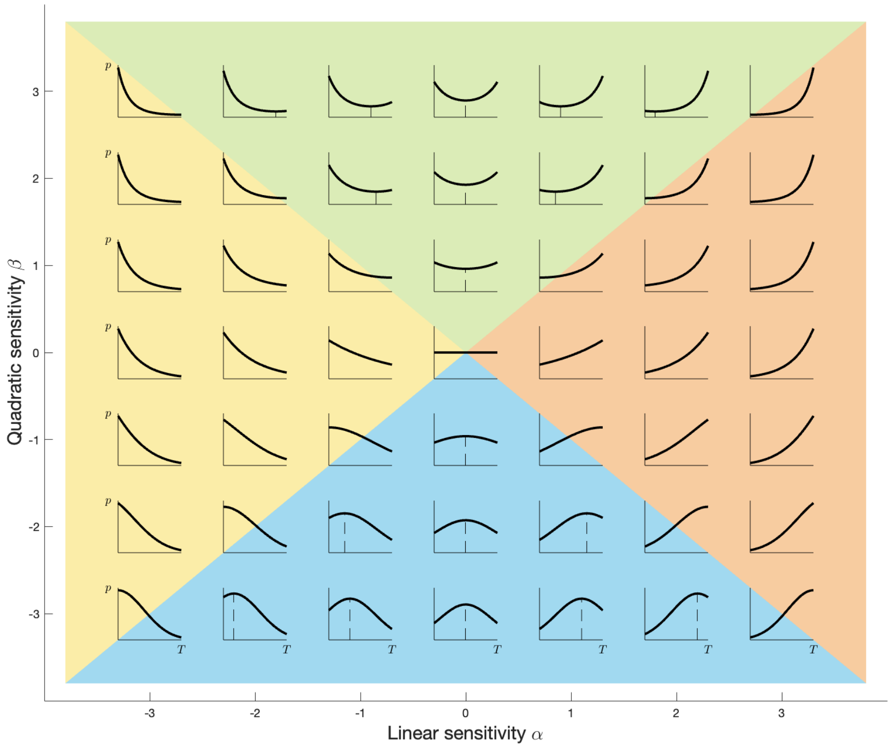


Figure S3: Graphical illustration of how the values of $\alpha_p$ (x-axis) and $\beta_p$ (y-axis) determine the shape of the thermal dependence of $p(T)$ (eq. S15). $\alpha_p$ and $\beta_p$ determine the sensitivity of the exponential function (and hence the thermal dependence) on the linear and quadratic parts of the exponent, respectively. A relative dominance of $\alpha_p$ leads to a monotonic shape, while $\beta_p$ relative dominance produces either hump- or U-shaped functions. Thus, four shapes emerge as defined by four regions, with smooth transitions between adjacent regions: monotonically increasing (red region), monotonically decreasing (yellow region), hump-shaped (blue region) or U-shaped (green region). Note: this is the same as Fig. 1 in the main text.

#### S2.2 Approximation example

Here, we demonstrate the conversion with an example, in which we transform a simple (and commonly used) form of the Arrhenius equation (eq. S19) into the Gaussian (eq. S20).

$$p_1(T) = p_{1,\mathrm{ref}} \exp\left(\frac{-E_a}{kT}\right) \tag{S19}$$

$$p_2(T) = p_{2,\mathrm{ref}} \exp\left(\alpha \frac{T - T_0}{T_w} + \beta \left(\frac{T - T_0}{T_w}\right)^2\right) \tag{S20}$$

In eq. (S19) we use notation specific to temperature parameterisations. Thus, $E_a$ refers to the activation energy, while $k$ is the Boltzmann constant. $p_{1,\mathrm{ref}}$ corresponds to the 'intercept', i.e., the value of the function for a given reference temperature (in this case $p_{1ref}$ is the value in the limit $T \to \infty$, see S2.2). Prior to converting eq. (S19) in to eq. (S20), the former can be log-transformed:

$$\ln\big(p_1(T)\big) = \ln\big(p_{1,\mathrm{ref}}\big) + \frac{-E_a}{kT} \tag{S21}$$

The conversion into the canonical form will now be based on the Taylor development of $\ln\big(p_1(T)\big)$ around $T \approx T_0$ up to the second order. First, we expand the reciprocal temperature terms from eq. (S21) in its Taylor series:

$$\frac{1}{T} = \frac{1}{T_0} + \frac{-1}{T_0^2}(T - T_0) + \frac{1}{T_0^3}(T - T_0)^2 + (\text{higher order terms})$$

Such that up to the second order in $(T - T_0)^2$ we have:

$$\frac{1}{T} = \frac{1}{T_0} + \frac{-1}{T_0^2}(T - T_0) + \frac{1}{T_0^3}(T - T_0)^2 \tag{S22}$$

Now we substitute the right-hand side of eq. (S22) for $\frac{1}{T}$ into the log-transformed Arrhenius (eq. S21):

$$\begin{aligned}
\ln(p_1(T)) &= \ln\big(p_{1,\mathrm{ref}}\big) + \frac{-E_a}{kT} \\
&= \ln\big(p_{1,\mathrm{ref}}\big) + \frac{-E_a}{k}\Big(\frac{1}{T_0} + \frac{-1}{T_0^2}(T - T_0) + \frac{1}{T_0^3}(T - T_0)^2\Big) \\
&= \ln\big(p_{1,\mathrm{ref}}\big) - \frac{E_a}{kT_0} + \frac{E_a}{kT_0^2}(T - T_0) - \frac{E_a}{kT_0^3}(T - T_0)^2
\end{aligned}$$

We can re-write this in the exponential form:

$$\begin{aligned}
p_1(T) &= \exp\Big(\ln(p_{1,\mathrm{ref}}) - \frac{E_a}{kT_0} + \frac{E_a}{kT_0^2}(T - T_0) - \frac{E_a}{kT_0^3}(T - T_0)^2\Big) \\
&= \exp\Big(\ln(p_{1,\mathrm{ref}}) - \frac{E_a}{kT_0}\Big)\ \exp\Big(\frac{E_a}{kT_0^2}(T - T_0) - \frac{E_a}{kT_0^3}(T - T_0)^2\Big) \\
&= p_{1,\mathrm{ref}}\ \exp\Big(\frac{E_a}{kT_0}\Big)\ \exp\Big(\frac{E_a}{kT_0^2}(T - T_0) - \frac{E_a}{kT_0^3}(T - T_0)^2\Big)
\end{aligned}$$

Thus, we can identify the transformations required to convert the parameters of eq. (S19) to the parameters $p_{2,\text{ref}}$, $\alpha$ and $\beta$ in the canonical form (eq. S20):

$$
\begin{aligned}
p_{2,\text{ref}} &= p_{1,\text{ref}} \exp\left(\frac{E_a}{kT_0}\right) \\
\frac{\alpha}{T_w} &= \frac{E_a}{kT_0^2} \qquad \text{or} \qquad \alpha = \frac{E_a}{kT_0^2} T_w \\
\frac{\beta}{T_w^2} &= \frac{E_a}{kT_0^3} \qquad \text{or} \qquad \beta = \frac{E_a}{kT_0^3} T_w^2
\end{aligned}
$$

Hence, the (empirically-determined) parameter values for eq. (S19) can be used to parameterise eq. (S20) as follows:

$$
\begin{aligned}
p_2(T) &= p_{1,\text{ref}} \exp\left(\frac{E_a}{kT_0}\right) \exp\left(\frac{E_a}{kT_0^2} T_w \frac{T - T_0}{T_w} + \frac{E_a}{kT_0^3} T_w^2 \left(\frac{T - T_0}{T_w}\right)^2\right) \\
&= p_{1,\text{ref}} \exp\left(\frac{E_a}{kT_0}\right) \exp\left(\frac{E_a}{kT_0^2}(T - T_0) + \frac{E_a}{kT_0^3}(T - T_0)^2\right)
\end{aligned}
$$

#### Note on interchangeable forms of the Arrhenius equation

In the literature, two different versions of the Arrhenius equation have been commonly used interchangeably (eqs. S23 and S24). In the approximation example above (eq. S19), we used the form without the term $T_{\text{ref}}$. However, certain parameterisations explicitly include the term $T_{\text{ref}}$.

$$
p(T) = p_{\text{ref}} \exp\left(\frac{E_a(T - T_{\text{ref}})}{kTT_{\text{ref}}}\right) = p_{\text{ref}} \exp\left(\frac{-E_a(T_{\text{ref}} - T)}{kTT_{\text{ref}}}\right) \tag{S23}
$$

$$
p(T) = p_\infty \exp\left(\frac{-E_a}{kT}\right) \tag{S24}
$$

Here, we demonstrate how eq. (S23) is equivalent to eq. (S24):

$$
\begin{aligned}
p(T) &= p_{\text{ref}} \exp\left(\frac{E_a(T - T_{\text{ref}})}{kTT_{\text{ref}}}\right) \\
&= p_{\text{ref}} \exp\left(\frac{E_a T}{kTT_{\text{ref}}}\right) \exp\left(\frac{-E_a T_{\text{ref}}}{kTT_{\text{ref}}}\right) \\
&= p_{\text{ref}} \exp\left(\frac{E_a}{kT_{\text{ref}}}\right) \exp\left(\frac{-E_a}{kT}\right)
\end{aligned}
$$

The first two terms are constants. We can define their product as $p_\infty$, such that $p_{\text{ref}}$ is the value of $p(T)$ at temperature $T_{\text{ref}}$, while $p_\infty$ is the value of $p(T)$ as temperature becomes very large, $T \to \infty$. Thus, we can switch from the form of eq. (S24) to eq. (S23).

#### S2.3 List of transformations

Here, we present the parameter transformations required to convert the functions we identified in the literature into the canonical form. We converted the following functions: the Arrhenius equation (eq. S25), exponential with linear and quadratic terms and reciprocal temperature (eq. S26), exponential with optimal temperature (eq. S27), exponential with linear and quadratic terms (eq. S28) and a non-exponential form with linear and square-root terms (eq. S29). In all conversions shown below we first present the function as used in the literature, followed by the necessary transformation of the three parameters in the canonical form (eq. S15), i.e., $p_0$, $\alpha$ and $\beta$.

We note some significant details in the following transformations. $T_0$ always refers to the temperature range mid-point, as defined in eq. (S17). $T_{\mathrm{ref}}$ and $T_{\mathrm{opt}}$ are the reference and optimal temperatures, respectively, in the original parameterisations. $k$ is the Boltzmann constant, $k = 8.62\ 10^{-5}\ \frac{\mathrm{eV}}{\mathrm{K}}$.

**Arrhenius equation**

$$p(T) = p_\infty \exp\left(-\frac{E_a}{kT}\right) \tag{S25}$$

$$\begin{aligned}
p_0 &= p_\infty \exp\left(-\frac{E_a}{kT_0}\right)\\
\alpha_p &= \frac{E_a}{kT_0}\frac{T_w}{T_0}\\
\beta_p &= -\frac{E_a}{kT_0}\left(\frac{T_w}{T_0}\right)^2
\end{aligned}$$

**Arrhenius quadratic form**

$$p(T) = p_{\mathrm{ref}} \exp\left(a\left(\frac{1}{kT}\right) + b\left(\frac{1}{kT}\right)^2\right) \tag{S26}$$

$$\begin{aligned}
p_0 &= p_{\mathrm{ref}} \exp\left(\frac{1}{kT_0}\left(a + \frac{b}{kT_0}\right)\right)\\
\alpha_p &= -\frac{1}{kT_0^2}\left(a + \frac{2b}{kT_0}\right)T_w\\
\beta_p &= -\frac{1}{kT_0^3}\left(a + \frac{3b}{kT_0}\right)T_w^2
\end{aligned}$$

**Based on optimal temperature**

$$p(T) = p_{\text{opt}} \, \exp\left(- \, a \, (T - T_{\text{opt}})^2\right) \tag{S27}$$

$$\begin{aligned}
p_0 &= p_{\text{opt}} \, \exp\left(- \, a \, (T_0 - T_{\text{opt}})^2\right) \\
\alpha_p &= -2a \, (T_0 - T_{\text{opt}}) T_w \\
\beta_p &= -a \, T_w^2
\end{aligned}$$

**Exponential-quadratic form**

$$p(T) = p_{ref} \, \exp\left(a \, (T - T_{\text{ref}}) + b \, (T - T_{\text{ref}})^2\right) \tag{S28}$$

$$\begin{aligned}
p_0 &= p_{ref} \, \exp\left(a \, (T_0 - T_{\text{ref}}) + b \, (T_0 - T_{\text{ref}})^2\right) \\
\alpha_p &= a \, T_w + 2b \, (T_0 - T_{\text{ref}}) \, T_w \\
\beta_p &= b \, T_w^2
\end{aligned}$$

**Linear-square-root form**

$$p(T) = a \, (T - T_{\text{lft}}) \sqrt{T_{\text{rgt}} - T} \tag{S29}$$

$$\begin{aligned}
p_0 &= a \, (T_0 - T_{\text{lft}}) \sqrt{T_{\text{rgt}} - T_0} \\
\alpha_p &= -\frac{T_w}{2} \frac{(T_0 - T_{\text{lft}}) - 2(T_{\text{rgt}} - T_0)}{(T_0 - T_{\text{lft}})(T_{\text{rgt}} - T_0)} \\
\beta_p &= -\left(\frac{T_w}{2}\right)^2 \frac{(T_0 - T_{\text{lft}})^2 + 2(T_{\text{rgt}} - T_0)^2}{(T_0 - T_{\text{lft}})^2 (T_{\text{rgt}} - T_0)^2}
\end{aligned}$$

#### S2.4 Application to data: parameter values

**Original parameterisations and parameter values**

Below we provide more details for the parameterisations in each study and the corresponding conversions. Specifically, we present the parameter values in the original studies and the corresponding $\alpha$s and $\beta$s of their conversions. We do not present original or converted intercept values, as these do not impact the shape of the warming-stability relationship, which is the focus of the study. We present the temperature values (for $T$, $T_{\text{opt}}$, etc.) in degrees Celcius, °C. However, to calculate the values of the temperature-dependent function, $p(T)$, one needs to work with temperature in Kelvin for the units to match up. Therefore, during the calculations one simply substitutes any $T$ in °C with $T + 273.15$ to the attain the value in Kelvin and perform the correct calculation. $k$ is the Boltzmann constant, $k = 8.62\ 10^{-5}\ \frac{\text{eV}}{\text{K}}$.

**Amarasekare (2015) (Amr)** Range 20−35°C, so that $T_w = 15$°C and $T_0 = 27.5$°C.

$$a(T) \propto \exp\Big(-\frac{(T - T_{\text{opt}})^2}{2s^2}\Big) \quad \text{with} \quad T_{\text{opt}} = 24°\text{C},\ s = 7.81°\text{C}$$
$$h(T) \propto \exp\Big(+\frac{(T - T_{\text{opt}})^2}{2s^2}\Big) \quad \text{with} \quad T_{\text{opt}} = 22°\text{C},\ s = 7.87°\text{C}$$
$$K(T) \propto \exp\Big(+\frac{(T - T_{\text{opt}})^2}{2s^2}\Big) \quad \text{with} \quad T_{\text{opt}} = 32°\text{C},\ s = 7.00°\text{C}$$

**Archer *et al.* (2019) (Arc1)** Range 5−25°C, so that $T_w = 20$°C and $T_0 = 15$°C.

$$a(T) \propto \exp\Big(\frac{-E_a(T_{\text{ref}} - T)}{kTT_{\text{ref}}}\Big) \quad \text{with} \quad E_a = 0.704\ \text{eV},\ T_{\text{ref}} = 10°\text{C}$$
$$h(T) = \text{constant}$$
$$K(T) \propto \exp\Big(\frac{-E_a(T_{\text{ref}} - T)}{kTT_{\text{ref}}}\Big) \quad \text{with} \quad E_a = 2.2295\ \text{eV},\ T_{\text{ref}} = 10°\text{C}$$

**Archer *et al.* (2019) (Arc2)** Range 5−25°C, so that $T_w = 20$°C and $T_0 = 15$°C.

$$a(T) \propto \exp\Big(\frac{-E_a(T_{\text{ref}} - T)}{kTT_{\text{ref}}}\Big) \quad \text{with} \quad E_a = 0.229\ \text{eV},\ T_{\text{ref}} = 10°\text{C}$$
$$h(T) = \text{constant}$$
$$K(T) \propto \exp\Big(\frac{-E_a(T_{\text{ref}} - T)}{kTT_{\text{ref}}}\Big) \quad \text{with} \quad E_a = 2.2295\ \text{eV},\ T_{\text{ref}} = 10°\text{C}$$

**Binzer *et al.* (2016) (Bin)** Range 0−40°C, so that $T_w = 40$°C and $T_0 = 20$°C.

$$a(T) \propto \exp\Big(\frac{-E_a(T_{\text{ref}} - T)}{kTT_{\text{ref}}}\Big) \quad \text{with} \quad E_a = 0.38\ \text{eV},\ T_{\text{ref}} = 20°\text{C}$$
$$h(T) \propto \exp\Big(\frac{-E_a(T_{\text{ref}} - T)}{kTT_{\text{ref}}}\Big) \quad \text{with} \quad E_a = -0.26\ \text{eV},\ T_{\text{ref}} = 20°\text{C}$$
$$K(T) \propto \exp\Big(\frac{-E_a(T_{\text{ref}} - T)}{kTT_{\text{ref}}}\Big) \quad \text{with} \quad E_a = -0.71\ \text{eV},\ T_{\text{ref}} = 20°\text{C}$$

**Fussmann *et al.* (2014) (Fus)** Range 0–40°C, so that $T_w = 40°C$ and $T_0 = 20°C$.

$$a(T) \propto \exp\left(\frac{-E_a(T_{\text{ref}} - T)}{kTT_{\text{ref}}}\right) \quad \text{with} \quad E_a = 0.581 \text{ eV},\ T_{\text{ref}} = 20°\text{C}$$
$$h(T) \propto \exp\left(\frac{-E_a(T_{\text{ref}} - T)}{kTT_{\text{ref}}}\right) \quad \text{with} \quad E_a = -0.467 \text{ eV},\ T_{\text{ref}} = 20°\text{C}$$
$$K(T) \propto \exp\left(\frac{-E_a(T_{\text{ref}} - T)}{kTT_{\text{ref}}}\right) \quad \text{with} \quad E_a = -0.772 \text{ eV},\ T_{\text{ref}} = 20°\text{C}$$

**Sentis *et al.* (2012) (Sen)** Range 13.9–32.8°C, so that $T_w = 18.9°C$ and $T_0 = 23.35°C$.

$$a(T) \propto \left(T - T_{\text{lft}}\right)\sqrt{T_{\text{rgt}} - T} \quad \text{with} \quad T_{\text{lft}} = 11.06°\text{C},\ T_{\text{rgt}} = 38.00°\text{C}$$
$$h(T) \propto \exp\left(\frac{c}{T}\right) \quad \text{with} \quad c = 44.76°\text{C},\ T \text{ expressed in } °\text{C}$$
$$K(T) = \text{constant}$$

**Uszko *et al.* (2017) (Usz)** Range 6–30°C, so that $T_w = 24°C$ and $T_0 = 18°C$.

$$a(T) \propto \exp\left(-\frac{(T - T_{\text{opt}})^2}{2s^2}\right) \quad \text{with} \quad T_{\text{opt}} = 22.85°\text{C},\ s = 9.4°\text{C}$$
$$h(T) \propto \exp\left(+\frac{(T - T_{\text{opt}})^2}{2s^2}\right) \quad \text{with} \quad T_{\text{opt}} = 20.95°\text{C},\ s = 7.2°\text{C}$$
$$K(T) = \text{constant}$$

**Vucic-Pestic *et al.* (2011) (Vuc1)** Range 5–30°C, so that $T_w = 25°C$ and $T_0 = 17.5°C$.

$$a(T) \propto \exp\left(\frac{-E_a(T_{\text{ref}} - T)}{kTT_{\text{ref}}}\right) \quad \text{with} \quad E_a = 0.37 \text{ eV},\ T_{\text{ref}} = 17.5°\text{C}$$
$$h(T) \propto \exp\left(\frac{-E_a(T_{\text{ref}} - T)}{kTT_{\text{ref}}}\right) \quad \text{with} \quad E_a = -0.24 \text{ eV},\ T_{\text{ref}} = 17.5°\text{C}$$
$$K(T) \propto \exp\left(\frac{-E_a(T_{\text{ref}} - T)}{kTT_{\text{ref}}}\right) \quad \text{with} \quad E_a = -0.36 \text{ eV},\ T_{\text{ref}} = 17.5°\text{C}$$

**Vucic-Pestic *et al.* (2011) (Vuc2)** Range 5–30°C, so that $T_w = 25°C$ and $T_0 = 17.5°C$.

$$a(T) \propto \exp\left(\frac{-E_a(T_{\text{ref}} - T)}{kTT_{\text{ref}}}\right) \quad \text{with} \quad E_a = -0.27 \text{ eV},\ T_{\text{ref}} = 17.5°\text{C}$$
$$h(T) \propto \exp\left(\frac{-E_a(T_{\text{ref}} - T)}{kTT_{\text{ref}}}\right) \quad \text{with} \quad E_a = -0.23 \text{ eV},\ T_{\text{ref}} = 17.5°\text{C}$$
$$K(T) \propto \exp\left(\frac{-E_a(T_{\text{ref}} - T)}{kTT_{\text{ref}}}\right) \quad \text{with} \quad E_a = -0.36 \text{ eV},\ T_{\text{ref}} = 17.5°\text{C}$$

**West & Post (2016) (Wes1)** Range 5−30°C, so that $T_w = 25$°C and $T_0 = 17.5$°C.

$$
\begin{aligned}
a(T) &\propto \exp\left(b\Big(\frac{-1}{kT}\Big) + q\Big(\frac{-1}{kT}\Big)^2\right) \qquad \text{with} \quad b = -100.3089 \text{ eV},\ q = -1.2789 \text{ eV}^2 \\
h(T) &\propto \exp\left(b\Big(\frac{-1}{kT}\Big) + q\Big(\frac{-1}{kT}\Big)^2\right) \qquad \text{with} \quad b = 60.233 \text{ eV},\ q = 0.7709 \text{ eV}^2 \\
K(T) &\propto \exp\left(b\Big(\frac{-1}{kT}\Big)\right) \qquad \text{with} \quad E_a = -0.30939
\end{aligned}
$$

**West & Post (2016) (Wes2)** Range 5−30°C, so that $T_w = 25$°C and $T_0 = 17.5$°C.

$$
\begin{aligned}
a(T) &\propto \exp\left(b\Big(\frac{-1}{kT}\Big) + q\Big(\frac{-1}{kT}\Big)^2\right) \qquad \text{with} \quad b = -39.5555 \text{ eV},\ q = -0.50128 \text{ eV}^2 \\
h(T) &\propto \exp\left(b\Big(\frac{-1}{kT}\Big) + q\Big(\frac{-1}{kT}\Big)^2\right) \qquad \text{with} \quad b = 31.0988 \text{ eV},\ q = 0.3936 \text{ eV}^2 \\
K(T) &\propto \exp\left(b\Big(\frac{-1}{kT}\Big)\right) \qquad \text{with} \quad b = -0.30939
\end{aligned}
$$

##### Converted parameter values

We used the conversions provided in equations (S25−S29) to compute the values for $\alpha$ and $\beta$ for the thermal dependencies of attack rate $a$, handling time $h$ and carrying capacity $K$ for each of the studies used. These values are listed in Table S1.

| study | attack rate $a$ | | | handling time $h$ | | | carrying capacity $K$ | | |
|---|---|---|---|---|---|---|---|---|---|
| | shape | $\alpha$ | $\beta$ | shape | $\alpha$ | $\beta$ | shape | $\alpha$ | $\beta$ |
| Amr | ∩ | −0.87 | −1.84 | ∪ | 1.33 | 1.82 | ∪ | −3.59 | 5.97 |
| Arc1 | ╱ | 1.97 | −0.14 | — | 0 | 0 | ╱ | 7.34 | 0 |
| Arc2 | ╱ | 0.64 | −0.04 | — | 0 | 0 | ╱ | 7.34 | 0 |
| Bin | ╱ | 2.1 | −0.28 | ╲ | −1.4 | 0.19 | ╲ | −3.8 | 0.52 |
| Fus | ╱ | 3.14 | −0.43 | ╲ | −2.52 | 0.34 | ╲ | −4.17 | 0.57 |
| Sen | ∩ | 0.9 | −1.60 | ╲ | −1.56 | 1.26 | — | 0 | 0 |
| Usz | ∩ | 1.32 | −3.26 | ∪ | −1.37 | 5.55 | — | 0 | 0 |
| Vuc1 | ╱ | 1.27 | −0.11 | ╲ | −1.37 | 0.07 | ╲ | −1.24 | 0.11 |
| Vuc2 | ╲ | −0.93 | 0.08 | ╲ | −0.79 | 0.07 | ╲ | −1.24 | 0.11 |
| Wes1 | ∩ | 6.12 | −15.60 | ∪ | −4.49 | 9.47 | ╲ | −1.06 | 0.09 |
| Wes2 | ∩ | 1.59 | −6.04 | ∪ | −1.10 | 4.73 | ╲ | −1.06 | 0.09 |

Table S1: Converted values for $\alpha$ and $\beta$ for each of the studies used.

#### S2.5 Application to data: graphical representation

Using the transformations provided in the previous section, we converted the empirically determined thermal dependencies for attack rate, handling time and carrying capacity from the literature (Fig. S4). In most cases, the converted (thin red curve) and original functions (thick grey curve) are identical. In very limited cases, the approximation deviates slightly from the original thermal dependence parameterisation. Given that the focus of this study is on the shape of the thermal dependence of stability (i.e., warming-stability relationship), we do not present the exact values of the intercept, i.e., $p_{1,\mathrm{ref}}$ and $p_{2,\mathrm{ref}}$. Instead, we eliminated this prefactor by normalising the studied parameter so that its maximum over the temperature range is equal to one. This normalisation allows us to compare the original and converted parameterisations (see Fig. S4).

These results would suggest that the conversion to the canonical form could be applied broadly in the literature. In this case, all temperature-dependent parameterisations could be represented in the form of eq. (S20), which would simplify cross-study and cross-system comparisons of the thermal dependence of specific vital rates (e.g., growth, metabolism), traits (e.g., attack rates) and environmental factors (e.g., carrying capacity).

**(a)** Attack rate

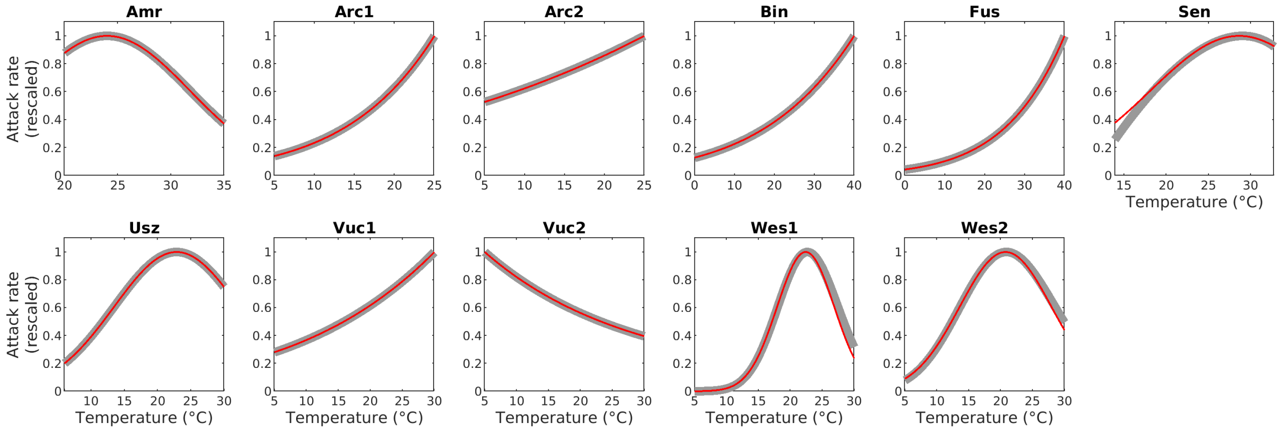


**(b)** Handling time

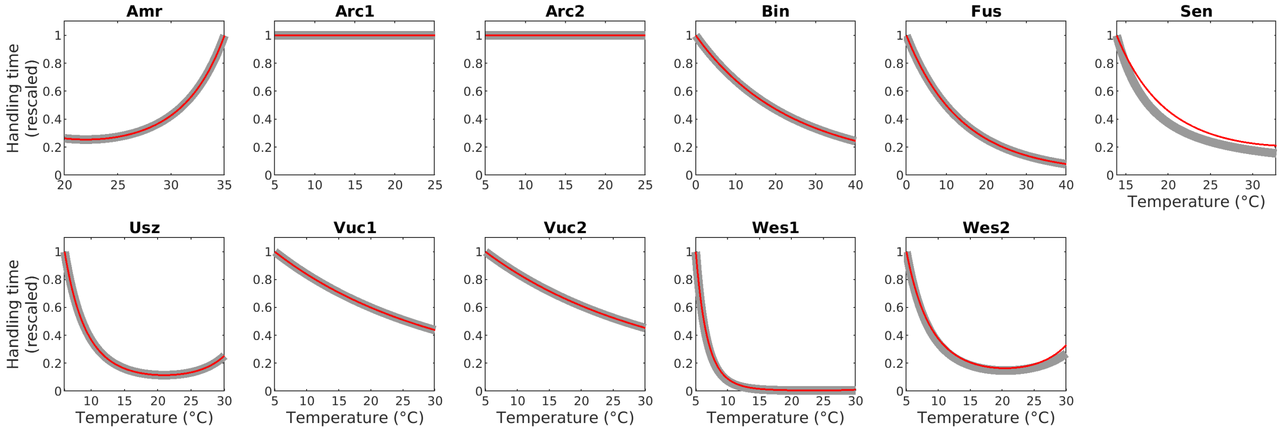


**(c)** Carrying capacity

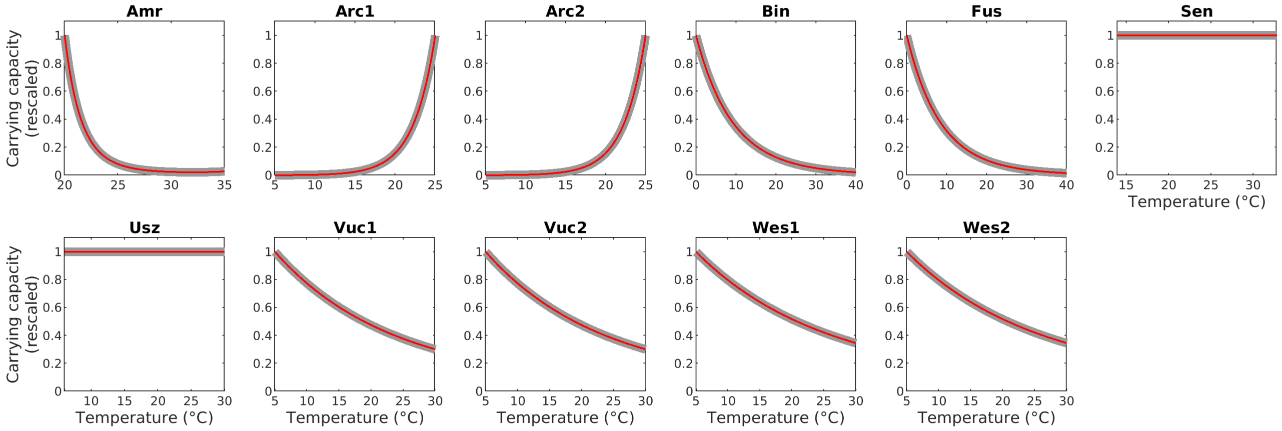


Figure S4: Comparison between original and converted temperature parameterisations. All these functions have been normalised by requiring that their maximum over the temperature range is equal to one. Thick grey line: original parameterisation; thin red line: converted parameterisation.

### Appendix S3: Graphical analysis of warming-stability relationships

#### S3.1 Contribution of individual parameters

$\varphi$ is the product of three parameters ($a$, $h$ and $K$), each of whose thermal dependence is represented by the same exponential function (*canonical* form developed in App. S2). Thus, the exponential thermal dependence of $\varphi$ can be represented on the $\alpha - \beta$ plane, with its $(x, y)$ coordinates determined by the respective sums $(\alpha_a + \alpha_h + \alpha_K,\ \beta_a + \beta_h + \beta_K)$. This means one can represent $\varphi$ in the $\alpha - \beta$ plane as the sum of three vectors, $\vec{a}$, $\vec{h}$ and $\vec{K}$. This representation illustrates the contribution of each parameter to the final position of $\varphi$ in the $\alpha - \beta$ plane, which determines the shape of $\varphi$ as a function of temperature and, hence, the warming-stability relationship.

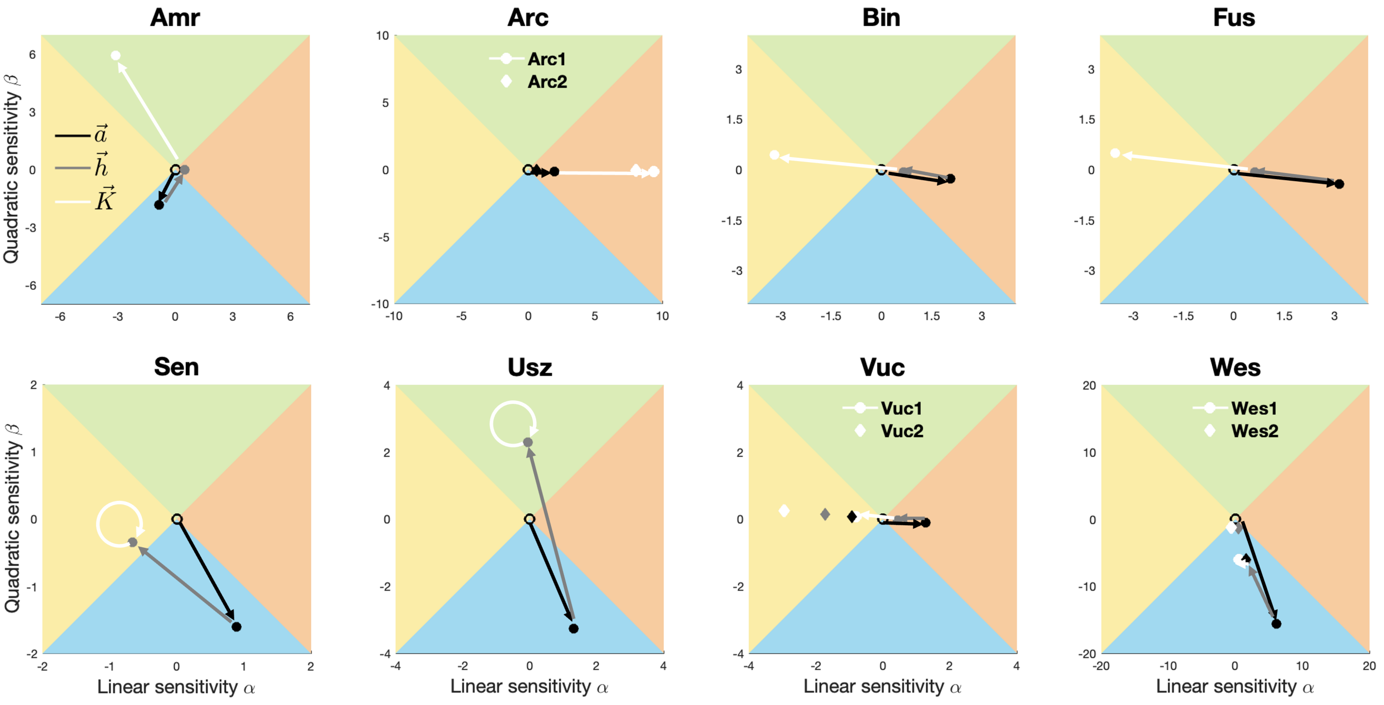


Figure S5: The 8 studies from the literature, whose data we mapped onto the $\alpha - \beta$ plane. For each study, we plot the three vectors, $\vec{a}$, $\vec{h}$ and $\vec{K}$, leading to the position of $\varphi$ on the plane.

#### S3.2 Sensitivity of warming-stability relationships

As we alluded to in the main text, different factors can contribute to uncertainty in the determination of the thermal dependence of any of the three parameters, $a$, $h$ or $K$. For instance, measurement errors or inaccurate assumptions could lead to (slight) deviations in the characterisation of the thermal dependence of a parameter and this can have significant implications for the overall warming-stability relationship. We demonstrate this *sensitivity* of the warming-stability outcome in two of the studies from our dataset.

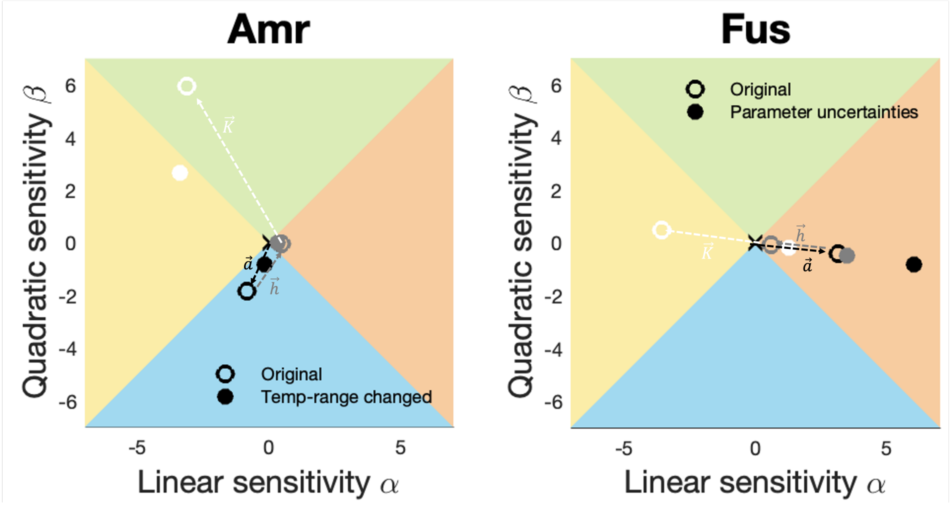


Figure S6: Sensitivity of the warming-stability relationship represented by changes in the position of $\varphi = (\alpha_\varphi, \beta_\varphi) = (\alpha_a + \alpha_h + \alpha_K,\ \beta_a + \beta_h + \beta_K)$. For both studies we present the outcome of the original parameterisation (open circles) and the outcome of an amended parameterisaiton (closed circles). For the latter, we shortened the temperature range over which the thermal dependence of the parameters was characterised for 'Amr', while for 'Fus' we used alternative parameter values for $a$ and $K$, based on the uncertainty (i.e., variance) in the original dataset.

In the first study (Fig. S6, Amr), the temperature range over which the thermal dependence of the parameters was characterised was altered from the original (Fig. S6, Amr, open circles) to a shortened range (Fig. S6, Amr, closed circles). In particular, the original measurements were conducted over a range of 20−35°C, while we artifially restricted this to 20−30°C. This change caused $(\alpha_\varphi, \beta_\varphi)$ to land in a different region (yellow instead of green), shifting the warming-stability relationship from hump-shaped (original) to monotonically stabilising (shortened temperature range).

The parameter values reported in the second study (Fig. S6, Fus) were derived from a meta-analysis of global invertebrate data (Fussmann *et al.*, 2014). The authors provided the mean values and the corresponding uncertainties (i.e., variance) of the activation energies of each parameter. We used the means as the 'original' values (Fig. S6, Fus, open circles). For the 'parameter uncertainties' (Fig. S6, Fus, closed circles) we used the values one standard deviation removed from the mean for $a$ and $K$. This uncertainty in the parameter values impacted the warming-stability outcome: from warming monotonically stabilising dynamics (yellow region), warming monotonically destabilises dynamics (red region).

In both examples, we induced changes in the thermal characterisation of the constituent parameters of $\varphi$, i.e., $a$, $h$ and $K$. Though artificial for the purposes of illustration, our changes were realistic. Determining the most appropriate temperature range over which to measure the thermal dependence of biological rates plays an important role in whether a rate will have a monotonic or unimodal thermal dependence. This figure highlights how choices made during the experimental design (whether imposed by technical constraints or other reasons) can impact the outcome of the experiment and, ultimately, what one would otherwise consider the 'correct' warming-stability relationship. Similarly, we demonstrated how measurement uncertainty in the parameter determination could smoothly shift the warming-stability relationship to a different, opposite even, outcome.
